# Sorafenib, a clinically approved kinase inhibitor attenuates *Streptococcus pneumoniae* pathogenesis *in vivo* by targeting serine/threonine kinase StkP

**DOI:** 10.1101/2025.08.14.670243

**Authors:** Joel Abraham, Aswathy C. Sagilkumar, Himani Dhyani, Charmi M. Panchal, Shaheena Aziz, M K Priyadatha, Aan Ruth, Keerthana Bhaskaran, Sivakumar Krishnankutty Chandrika, Rosemol Varghese, Ayyanraj Neeravi, Balaji Veeraraghavan, Sandhya Ganesan, Karthik Subramanian

**Author notes:** Correspondence to lead contact, Dr. Karthik Subramanian, Faculty Scientist E1, Host-Pathogen Laboratory, BRIC-Rajiv Gandhi Centre for Biotechnology (BRIC-RGCB), Thiruvananthapuram, 695014, India.

## Abstract

*Streptococcus pneumoniae* is a respiratory commensal bacterium responsible for over one million annual fatalities globally, particularly among children under five years of age. The rapid emergence of macrolide-resistant strains led the WHO in 2024 to designate *S. pneumoniae* as a priority pathogen, underscoring the need for alternative strategies such as anti-virulence therapy. Here, we repurposed the FDA-approved cancer drug, sorafenib identified by *in silico* screening of compounds targeting the bacterial Serine/Threonine kinase protein StkP, an essential regulator of cell division and peptidoglycan synthesis conserved across many bacterial pathogens. Sorafenib interacts with the StkP-kinase domain and exhibited broad-spectrum activity against diverse pneumococcal serotypes including multi-drug-resistant clinical isolates. Ectopic expression of StkP in both wild-type and isogenic mutant strains conferred partial resistance to sorafenib, confirming on-target activity. Scanning electron microscopy revealed aberrant cell-wall morphology, and differential viability staining demonstrated increased membrane permeability. Consistently, sorafenib-treated bacteria showed significantly higher complement C3 deposition and consequent killing in human serum. In human lung epithelial cells, sorafenib reduced bacterial adherence and invasion without detectable host cytotoxicity. Serial passaging at sub-microbicidal concentrations indicated a low propensity for resistance development *in vitro*. *In vivo*, sorafenib administration reduced mortality and lung bacterial load in a mouse model of pneumonia. Taken together, our data identify StkP as target of sorafenib in *S. pneumoniae* and justify its continued preclinical development for therapeutic intervention.

## Introduction

*Streptococcus pneumoniae* (pneumococcus) is a Gram-positive human pathogen responsible for many invasive (pneumonia, meningitis, sepsis) and non-invasive (otitis media) diseases, causing an estimated 1.6 million deaths annually (Narciso *et al*, 2024) (Weiser et al., 2018). The global burden of pneumococcal infections is substantial, with an estimated 1.6 million deaths annually, particularly affecting young children, the elderly, and immunocompromised individuals (World Health Organization, 2019). Notably, *S. pneumoniae* was the highest causative of mortality among young children post-neonatal to age 4 years (Collaborators, 2022). The clinical burden is exacerbated by the rapidly rising antimicrobial resistance that pose significant challenges to effective treatment and disease management (Cherazard *et al*, 2017). Over the past few decades, the emergence and spread of multidrug-resistant pneumococcal strains has been attributed to the widespread and often indiscriminate use of antibiotics, as well as the bacterium’s remarkable ability to acquire and disseminate resistance genes through horizontal gene transfer (Hakenbeck *et al*, 2012). Resistance to currently used antibiotic classes such as beta-lactams, macrolides, and tetracycline has been reported and multi-drug-resistant strains are emerging worldwide (Gergova *et al*, 2024; Sharew *et al*, 2021). In 2024, the WHO classified the macrolide-resistant *S. pneumoniae* as one of twelve priority pathogens in urgent need of new therapeutics.

Anti-virulence strategies, which disarm pathogens without directly inhibiting growth offer a promising alternative that may impose less selective pressure for resistance (Johnson & Abramovitch, 2017) (Rasko & Sperandio, 2010). Several virulence factors contribute to the pathogenicity of *S. pneumoniae*, including the polysaccharide capsule, surface proteins, and various proteins involved in host-pathogen interactions (Narciso *et al*., 2024). Among these, the serine/threonine kinase protein, StkP has garnered significant attention as a potential therapeutic target. StkP is a key regulator of pneumococcal physiology and virulence, playing crucial roles in cell division, cell wall biosynthesis, competence and stress response (Beilharz et al., 2012). In *S. pneumoniae*, StkP has been shown to phosphorylate multiple substrates, including cell division proteins (FtsZ, DivIVA), cell wall biosynthesis enzymes (MurC, GlmM), and transcriptional regulators (RitR) (Fleurie *et al*, 2012) (Falk & Weisblum, 2013). Through these interactions, StkP modulates various cellular processes that are critical for pneumococcal survival and virulence. Genetic disruption of StkP attenuates virulence in animal models, impairs cell division, diminishes biofilm formation, and increases sensitivity to environmental stressors, all of which contribute to decreased virulence (Echenique *et al*, 2004) (Beilharz *et al*, 2012). Several characteristics make StkP an attractive target for therapeutic intervention. Firstly, its conservation across pneumococcal strains and presence of homologous protein, PknB in *Staphylococcus aureus* (Huemer et al, 2023) and *Mycobacterium tuberculosis* (Chawla et al, 2014) suggests that targeting StkP could provide broad-spectrum activity against many priority pathogens. Secondly, the multifaceted role of StkP in pneumococcal physiology implies that its inhibition could simultaneously affect multiple virulence mechanisms, potentially enhancing therapeutic efficacy.

Here, we screened 38 host-targeted kinase inhibitors against the pneumococcal StkP kinase domain and identified sorafenib as a lead compound, which is currently FDA-approved drug for hepatocellular carcinoma (Llovet *et al*, 2008). Although sorafenib is clinically used to inhibit VEGFR, PDGFR, and RAF kinases in treatment of hepatocellular carcinoma, a recent report noted its activity against methicillin-resistant *S. aureus* via menaquinone pathway inhibition, without a defined kinase target (Le et al, 2020). Before developing sorafenib or its derivatives for antimicrobial therapy, it is crucial to elucidate the precise mechanisms through which it demonstrates antimicrobial activity against different pathogenic bacteria. *Streptococcus pneumoniae* is a unique pathogen due to the presence of over 100 different capsular serotypes and expresses the eukaryotic-type bacterial kinase, StkP, which is a key virulence factor. To elucidate sorafenib’s mechanism against pneumococcus, we used molecular dynamics simulations to map its binding to the StkP active site in competition with the natural substrate DivIVA and identified key interacting residues. We then characterized antimicrobial activity through kinetic growth assays, membrane permeability studies, complement-mediated serum killing, and epithelial cell invasion assays. Efficacy was tested against multidrug-resistant clinical isolates, and on-target engagement was confirmed by partially rescuing sorafenib inhibition through Zn-inducible expression of StkP in both wild-type and knockout backgrounds. Serial passaging over 16 generations at sub-microbicidal concentrations of sorafenib revealed no significant rise in the minimal inhibitory concentration, underscoring its low propensity for resistance development. The efficacy of sorafenib was further validated in a mouse oropharyngeal infection model.

## Results

### Sorafenib binds to the catalytic site of pneumococcal StkP *in silico*

The pneumococcal eukaryotic-type kinase StkP phosphorylates downstream substrate proteins like DivIVA (Fleurie *et al*, 2014; Novakova *et al*, 2010), FtsZ (Giefing *et al*, 2010), MapZ (LocZ) (Holeckova *et al*, 2014), MacP (Fenton *et al*, 2018), KhpB (Jag/EloR) (Ulrych *et al*, 2016) (Stamsas *et al*, 2017) that are collectively involved in peptidoglycan regulation for the cell wall homeostasis (Beilharz *et al*., 2012). The N-terminal region of StkP protein consists of a catalytically active, cytosolic kinase domain (hereafter referred as StkP-KD) spanning to 273 amino acids, followed by a juxtamembrane region (274-344), transmembrane domain (345-363), and four PASTA domains (364-434, 435-504, 505-577, 578-651) (**Fig. S1A**). StkP-KD consists of a catalytic loop, activation loop, P-loop, and P+1 loop which are involved in the phosphorylation activity of StkP (Righino *et al*, 2018). Since there is no available crystal structure for StkP or the kinase domain, the three-dimensional structure was predicted using the neural network-based model, AlphaFold (Jumper *et al*, 2021) (**Fig. S1B**) and the generated structure was validated using Ramachandran plot analysis (**Fig. S1C**). Additionally, the StkP-KD structure was superimposed with the available crystal structures of homologous kinase proteins, PknB (*S. aureus)*, PknA and PknB (*M. tuberculosis)* with 47.84, 42.17 and 41.26% sequence identities respectively (**Fig. S1D**). The residues in the catalytic loop, Val-133 to Asn-141 and ATP-binding P-loop of StkP-KD were highly conserved in all three bacterial species. Pneumococcal StkP-KD showed a high structural similarity with that of *Staphylococcal* and *Mycobacterial* PknA and PknB proteins with root-mean square deviation (RMSD) values of 1.053, 1.316 and 1.006 Å respectively (**Fig. S1E**).

Next, we tested the similarity of pneumococcal StkP-KD with human kinases at the sequence and structural levels. We found that there was only 30% overall sequence similarity (**Fig. S2A**), but catalytic residues, His-134, Arg-135, Asp-136 and Leu-137 in the catalytic loop as well as glycine rich P-loop in the ATP binding site were conserved (**Fig. S2B**). Structural homology modelling revealed the conservation of β-strand-loop-β-strand motif in ATP binding pocket and catalytic loop (**Fig. S2C**). This prompted us to hypothesize that human kinase inhibitors could be repurposed against pneumococcal StkP.

We screened 38 host-targeted kinase inhibitors (**Table S1**) that were previously reported to show activity against liver-stage malaria infection (Arang *et al*, 2017) against the pneumococcal StkP. Molecular docking analysis of the molecules with StkP-KD was performed using the Glide extra-precision scoring function on the Schrodinger suite (Friesner *et al*, 2006). The top two compounds with the best docking scores were Dasatinib and Sorafenib with -8.304 and -8.279 kcal/mol respectively were chosen for experimental validation (**Table S2**). Growth inhibition analysis showed that sorafenib completely inhibited the growth of *S. pneumoniae* TIGR4 strain as compared to dasatinib at 10 µm. (**Fig. S2D**).

Docking studies of StkP-KD with sorafenib showed stable binding to the catalytic cleft of the protein (**Fig. 1A**). A 200 ns molecular dynamics simulation of the docked complex showed consistent hydrogen bonding and hydrophobic bonding interactions with the catalytic residues, Arg-135 and Asp-136 (**Figs. 1B and S3A, B**). Furthermore, RMSD analysis of the complex over the 200 ns molecular dynamics trajectory exhibited minimal deviation, corroborating the formation of a stable complex. (**Fig. 1C and Videos S1, S2**). To verify the involvement of the catalytic loop residues, Arg-135 and Asp-136 in binding to sorafenib, the simulation was performed with both residues mutated to alanine. The mutations abolished interaction with the StkP-KD catalytic loop (**Figs. S3C-E**), resulting in ∼2.5-fold higher RMSD value for sorafenib (**Fig. 1D**), which eventually caused the sorafenib to drift away from the binding pocket (**Video S3**). DivIVA is a substrate of StkP which is phosphorylated at the Thr-201 position (Fleurie *et al*., 2012), and plays vital role in proper septum formation and nucleoid segregation during cell division. Due to the absence of a crystal structure, the structure of DivIVA peptide was modeled using AlphaFold and a nine amino acid peptide spanning the Thr-201 was used in competition with sorafenib to test if it can replace sorafenib from the binding pocket of StkP-KD. DivIVA peptide was placed on a near to the catalytic area of StkP-KD along with sorafenib. A 200 ns simulation showed that the peptide could not replace sorafenib in the binding pocket and eventually moved away from the active site (**Videos S4 and S5**). While sorafenib showed persistent interactions with StkP catalytic residues in the presence of DivIVA (**Figs. S4A, B**), DivIVA did not interact with the catalytic site in the presence of sorafenib (**Fig. S4C, D)**. Moreover, both StkP-KD and sorafenib showed a stable RMSD throughout the simulation period, whereas DivIVA displayed high RMSD fluctuations at several points of simulation period with ∼4-fold final RMSD value compared to sorafenib indicating unstable interactions in the presence of sorafenib (**Figs. 1E, F**). We also performed Molecular Mechanics with Generalized Born and Surface Area solvation (MM/GBSA) analysis to calculate the free energy of the binding of sorafenib in simulation systems containing sorafenib alone and in the presence of DivIVA or StkP-KD R135A/D136A double mutant. MM/GBSA analysis showed lower free energy of sorafenib with StkP as compared to DivIVA (**Fig. 1G)**. Besides, mutation of Arg-135 and Asp-136 residues to alanine also increased free energy of binding between sorafenib and StkP-KD. Taken together, our data shows that sorafenib binds to the catalytic site of pneumococcal StkP-KD in a highly stable manner, interacting with the catalytic residues, even in the presence of a natural StkP substrate. Together, these findings identify the catalytic domain of pneumococcal StkP as a structurally druggable target and demonstrate that sorafenib binds stably to its active site by engaging with key catalytic loop residues.

**FIGURE 1.**
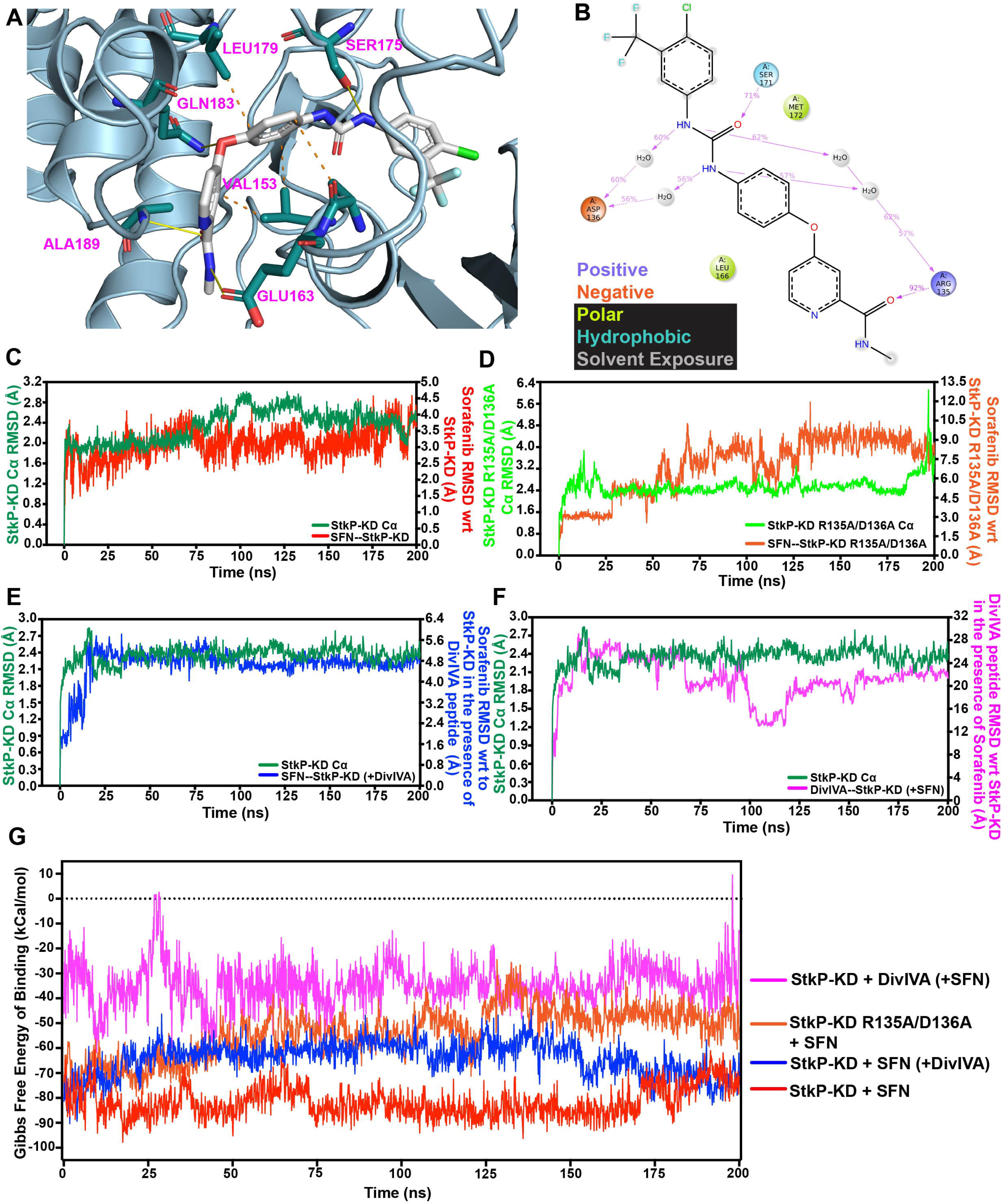
Sorafenib binds to the catalytic site of pneumococcal StkP *in silico*. **(A).** Docked pose of sorafenib (SFN) at the catalytic site of pneumococcal StkP-Kinase Domain (StkP-KD), showing the major amino acids involved in initial hydrogen bonding (yellow straight lines) and hydrophobic (orange dotted lines) interactions. **(B).** Contact diagram showing the interactions between SFN and StkP-KD during a 200 ns molecular dynamic simulation. Amino acids that interacted for more than 30% of the total time are shown here. **(C-D).** RMSD analysis of the structural fluctuations of (**C**) StkP-KD wild-type and (**D**) StkP-KD R135A/D136A α-carbon backbone - and SFN with respect to the proteins. **(E-F).** RMSD analysis of (**E**). StkP-KD α-carbon backbone and SFN with respect to StkP-KD in the presence of DivIVA peptide and **(F).** StkP-KD α-carbon backbone and DivIVA peptide with respect to StkP-KD in the presence of SFN. **(G).** Gibb’s free energy of binding of SFN to wild-type StkP-KD and StkP-KD R135A/D136A showing the importance of Arg 135 and Asp 136 during the interaction with SFN. The relative binding energies of SFN and DivIVA peptide to StkP-KD in a triple molecule simulation is shown indicating lower binding energy for StkP-SFN compared to StkP-DivIVA peptide in the presence of SFN.

### Sorafenib effectively counters multidrug resistant pneumococci by targeting StkP

Next, we wanted to study the potential antimicrobial effects of sorafenib *in vitro*. Pneumococcal strains, TIGR4 (serotype 4) and D39 (serotype 2) were grown in presence of sorafenib at concentrations ranging between 0.2-5 μM. Equivalent concentrations of the solvent, DMSO was used as the control. We found that sorafenib induced a dose-dependent inhibition of bacterial growth with minimum inhibitory concentration (MIC) value of 2.5 μM. (**Figs. 2A, B**). Further, we tested the activity of sorafenib against six clinical pneumococcal strains, out of which five isolates were resistant to the commonly used macrolide, erythromycin with high MIC value of >128 μg/mL (**Table 1**). Our results showed that sorafenib was effective against all the strains tested irrespective of penicillin and erythromycin resistance. Interestingly, sorafenib showed ∼4-fold lower MIC when compared to erythromycin in two clinical isolates, GLO00007 and SP676 with high erythromycin resistance and intermediate penicillin resistance (**Table 1 and Fig. 2C**).

**FIGURE 2.**
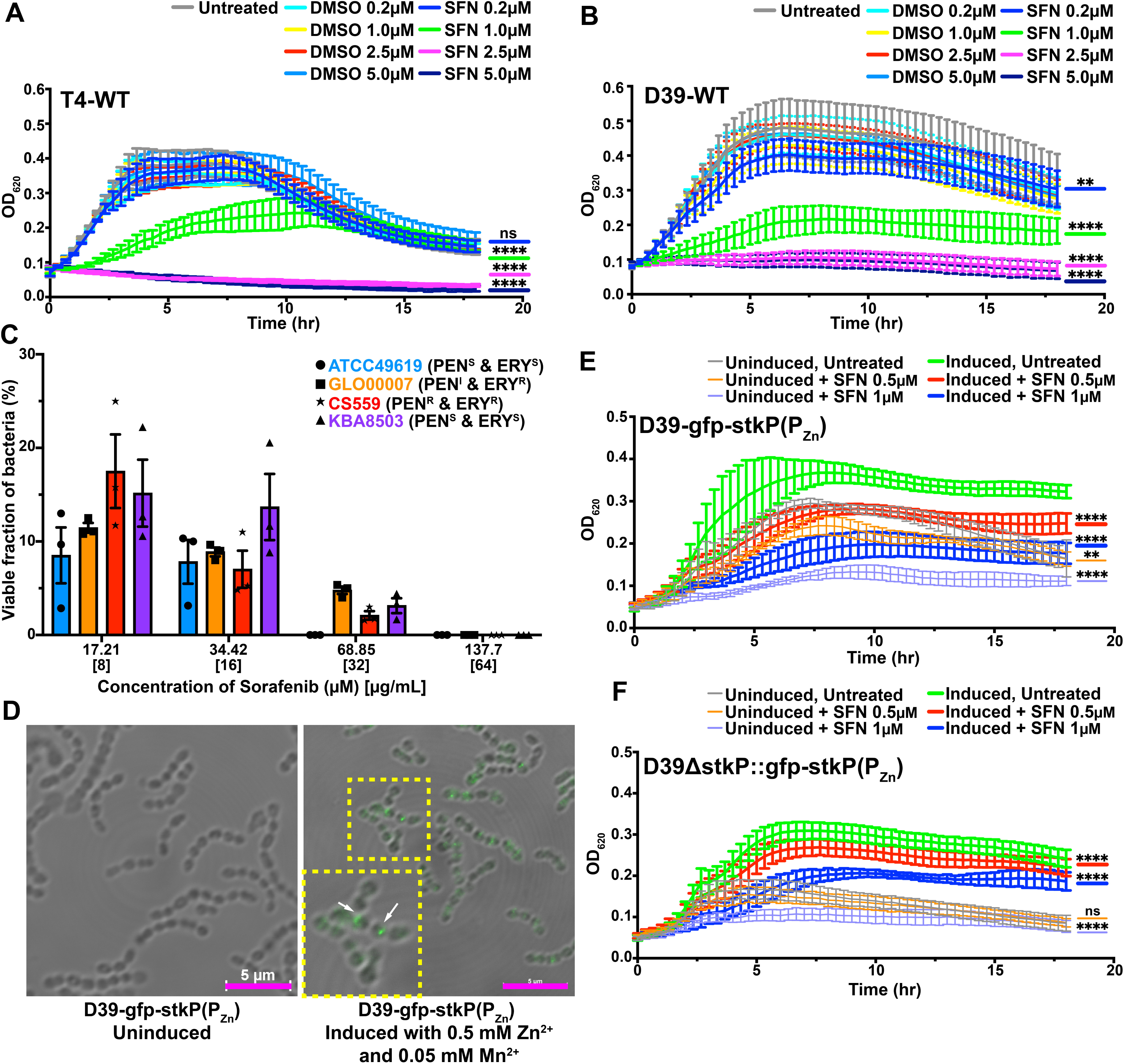
Sorafenib effectively counters multidrug resistant pneumococci by targeting StkP. **(A-B).** Growth kinetics of wild-type *S. pneumoniae* strains, (**A**) TIGR4 (T4) and (**B**) D39, showing dose-dependent inhibition of growth with increasing concentrations of sorafenib (SFN). Equimolar concentrations of DMSO (solvent) and untreated bacteria served as negative controls. ** indicates p ≤ 0.01 and **** indicates p ≤ 0.0001 relative to DMSO; ns denotes non-significance by Mann-Whitney test. **(C).** Percentage viability of erythromycin (ERY) and penicillin (PEN) resistant and sensitive clinical strains of *S. pneumoniae* upon treatment with different concentrations of SFN. R denotes resistant, S-sensitive and I-intermediate following the CLSI criteria. **(D).** Confocal microscopy images of D39-gfp-stkP(P_Zn_) strain showing the localization of GFP-StkP at the division septum upon induction with 0.5 mM ZnCl_2_ and 0.05 mM MnCl_2_ compared to uninduced condition. Inset shows magnification of selected region. Scale bars, 5 µm. **(E, F).** Growth kinetics of D39-gfp-stkP(P_Zn_) and D39ΔstkP::gfp-stkP(P_Zn_) respectively, upon treatment with 0.5 and 1 µM of SFN in both uninduced and induced conditions. ** indicates p ≤ 0.01 and **** indicates p ≤ 0.0001 comparing uninduced/induced SFN treated bacteria to the respective untreated bacteria; ns denotes non-significance by Mann-Whitney test. All data are representative of mean ± SEM from three independent experiments.

**Table 1:** MIC values of sorafenib against multi-drug-resistant clinical strains of pneumococci, S, susceptible; I, intermediate; R, resistant following the CLSI criteria.

| **ID** | **Child/Adult** | **Source** | **Invasive/Non-invasive** | **Diagnosis** | **Serotype** | **Penicillin MIC (µg/mL)** | **Erythromycin MIC (µg/mL)** | **Sorafenib MIC (µg/mL) [µM]** | **DMSO MIC (µg/mL)** |
| --- | --- | --- | --- | --- | --- | --- | --- | --- | --- |
| **GLO00007** | Child | Pleural fluid | invasive | Pneumonia | 19A | **4 (I)** | **>128 (R)** | 32<br>[68.85] | >128 |
| **SP676** | Child | sputum | non invasive | Respiratory Tract Infection | 19F | **4 (I)** | **>128 (R)** | 32<br>[68.85] | >128 |
| **SP2782** | Adult | sputum | non invasive | Respiratory Tract Infection | 19F | **2 (S)** | **>128 (R)** | 32<br>[68.85] | >128 |
| **CS559** | Adult | CSF | invasive | Meningitis | 19F | **2 (R)** | **8 (R)** | 32<br>[68.85] | >128 |
| **311104** | child | nasopharyngeal swab | non invasive | carriage | 19F | **2 (S)** | **8 (R)** | 32<br>[68.85] | >128 |
| **KBA8503** | Adult | Blood | invasive | Bacteraemia | 6C | **0.12 (S)** | **0.25 (S)** | 32<br>[68.85] | >128 |

To investigate whether StkP is a potential target of sorafenib, we used genetically engineered pneumococcal D39 strain to express the N-terminal GFP-tagged StkP (GFP-StkP) at the dispensable bgaA locus under a zinc-inducible P_czcD_ promoter (P_Zn_) in the wild-type (D39-gfp-stkP(P_Zn_)) as well as the knockout strain (D39ΔstkP::gfp-stkP(P_Zn_)). The gradient induction of GFP-StkP expression in the D39-gfp-stkP(P_Zn_) strain using ZnCl_2_ and MgCl_2_ was validated by flow cytometry (**Fig. S5A**) and western blotting (**Fig. S5B**). The septal localization of GFP-StkP in the bacterial cells was confirmed by confocal microscopy (**Fig. 2D**) as previously reported (Beilharz *et al*., 2012). The genomic integration of ectopic StkP in the wild-type D39 strain, D39-gfp-stkP(P_Zn_) and the knockout strain D39ΔstkP::gfp-stkP(P_Zn_) was validated by PCR (**Fig. S5C**) and protein expression confirmed by flow cytometry (**Fig. S5D)**. The septal localization of the GFP-StkP in the complemented strain was validated by confocal microscopy (**Fig. S5E**). Growth kinetic assay showed that ectopic expression of StkP in D39-gfp-stkP(P_Zn_) strain and the complemented knockout strain, D39ΔstkP::gfp-stkP(P_Zn_) was able to be partly rescue growth in a dose-dependent manner to 0.5-1 µM of sorafenib (**Figs. 2E and 2F**). Upon StkP induction, D39-gfp-stkP(P_Zn_) showed ∼1.5-fold higher growth in the presence of 1 µM sorafenib with 55% growth recovery relative to untreated compared to ∼46% in uninduced condition, suggesting that StkP is a target of sorafenib **(Fig. 2E).** No dose-responsiveness was observed with the solvent, DMSO relative to untreated **(Fig.S5F)**. Similarly, the complemented knockout strain, D39ΔstkP::gfp-stkP(P_Zn_) showed a ∼1.8-fold higher growth upon StkP induction compared to uninduced and was comparable to the untreated (**Fig. 2F and S5G**).

### Sorafenib blocks the kinase activity of StkP

To investigate the effect of sorafenib on the kinase activity of StkP, we analyzed the global phosphorylation profile in *S. pneumoniae* D39 and TIGR4 strains following treatment with sorafenib. Immunodetection using a phospho-threonine-specific (anti-pThr) antibody revealed a marked reduction in the phosphorylation levels of several bacterial proteins compared to DMSO-treated controls (**Figs. 3A and S6A, C**). Enolase was used as the loading control (**Fig. 3B and S6B, D**) and used for densitometry analysis (**Fig. 3C**). Mass-spectrometry identification of the proteins below 37 kDa that showed downregulation, revealed proteins, DivIVA and GpsB that are substrates of StkP and involved in bacterial cell division (**Table 2**). Next, we recombinantly expressed and purified His-tagged StkP kinase domain (StkP-KD) in *E. coli* BL21(DE3) cells (**Fig. S6E and F**) to directly assess the impact of sorafenib on StkP autophosphorylation using an *in vitro* kinase assay. The autophosphorylation activity of StkP-KD was examined by incubating the purified protein with ATP in kinase buffer. As expected, StkP-KD exhibited a dose-dependent increase in phosphorylation with increasing ATP concentrations (**Fig. S6G, H**). Notably, incubation with sorafenib, but not with equivalent concentrations of the solvent DMSO, led to a dose-dependent reduction in StkP-KD phosphorylation (**Fig. S6I-K**), confirming the direct inhibitory effect of sorafenib on StkP kinase activity. Since the purified StkP-KD was already showing a basal level of phosphorylation upon expression in *E. coli*, we directly examined the autophosphorylation of StkP-KD in transformed *E. coli,* cultures grown without IPTG induction to permit only basal-level expression and grown in the presence of sorafenib. Our results demonstrated that sorafenib, but not the solvent DMSO, caused a significant and dose-dependent reduction in StkP phosphorylation, relative to untreated (**Fig. 3D and E**). The total cell lysate was used as the loading control (**Fig. S6L**). Given that *E. coli* lacks eukaryotic-like Ser/Thr kinases and is therefore not inherently involved in the regulation of cell division (Macek *et al*, 2007), we hypothesized that sorafenib would not affect bacterial growth in this heterologous expression system. Consistent with this, the growth kinetics of *E. coli* expressing StkP-KD remained unaltered in the presence of sorafenib, supporting the specificity of its inhibitory effect (**Fig. S6M**). Together, these results establish sorafenib as a potent inhibitor of *S. pneumoniae* by targeting the Ser/Thr kinase, StkP.

**FIGURE 3:**
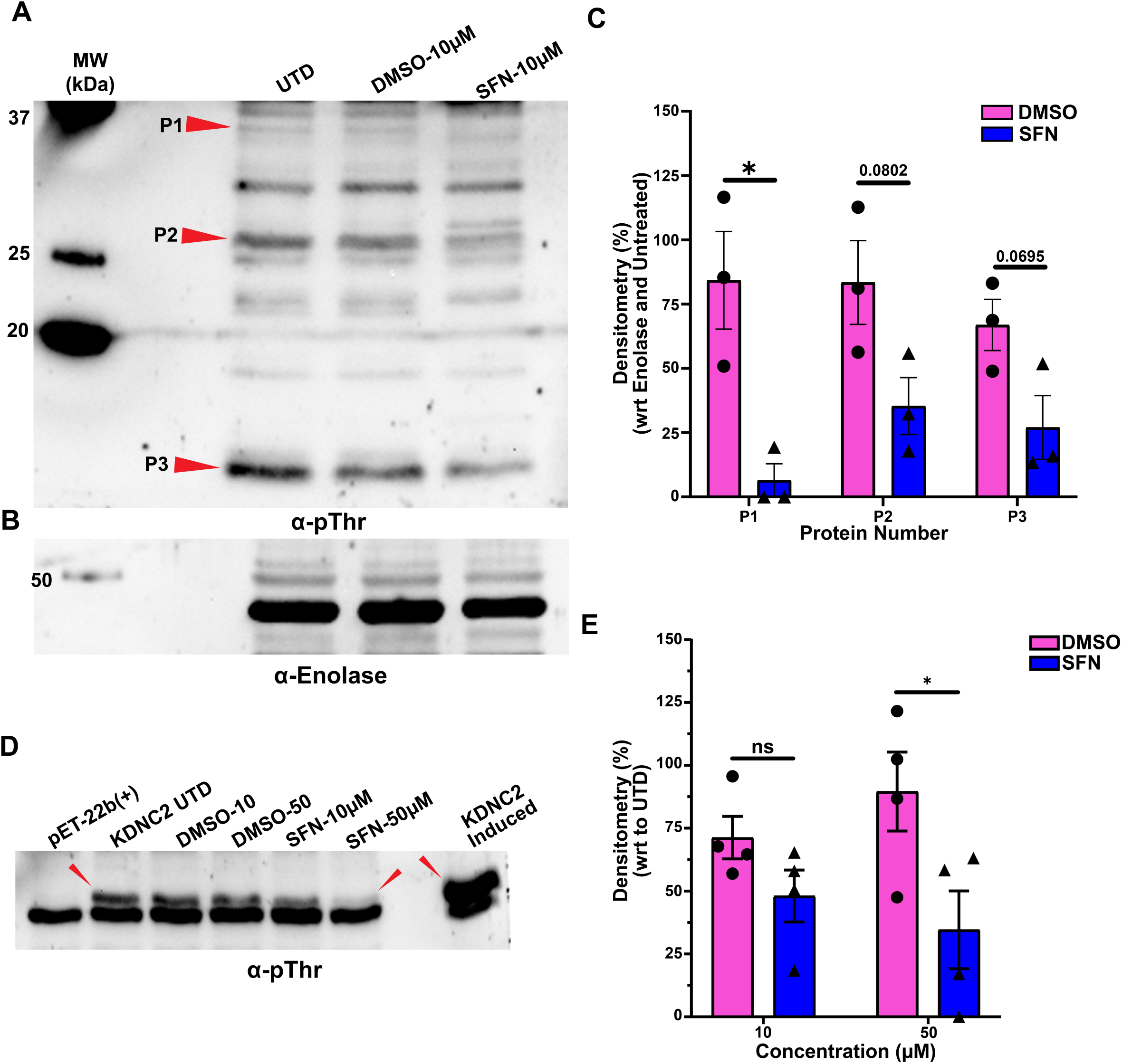
Sorafenib blocks the kinase activity of StkP. **(A, B)** Western blots probed with (**A**) pThr specific antibody and (**B**) enolase (loading control) showing the downregulation of phosphorylated proteins in *S. pneumoniae* D39 strain upon treatment with 10 μM sorafenib (SFN). Untreated (UTD) and equivalent amount of DMSO treated bacteria were used as the controls. (**C**). Densitometry analysis of downregulated bands normalized to enolase and untreated control from three independent experiments. **(D).** Western blot probed with pThr specific antibody showing the dose-dependent downregulation of phosphorylation of ectopically expressed pneumococcal StkP-KD in *E. coli* BL21(DE3) cells without induction and upon treatment with SFN. Non-transformed BL21(DE3) cells and transformed cells with StkP-KD expression induced using 0.50 mM IPTG were used as controls. Arrow indicates StkP-KD band. **(E)**. Densitometry analysis for panel D, normalized to untreated bacteria. * indicates p ≤ 0.05 and ns denotes non-significance by Welch’s t-test. Data are representative of four independent experiments.

**Table 2:** List of proteins in selected bands below 37 kDa in Figure 3A identified from Mass Spectrometry.

| **UNIPROT ID** | **Protein Name** | **Molecular Weight (kDa)** | **Gene Ontologies** |
| --- | --- | --- | --- |
| A0A062WNB1 | Cell division protein SepF | 20.6 | Division Septum Assembly [0000917], FtsZ-Dependent Cytokinesis [0043093] |
| A0A062WWL4 | Glutamate racemase | 29.1 | Cell Wall Organization [0071555], Peptidoglycan Biosynthetic Process [0009252], Regulation of Cell Shape [0008360] |
| A0A064C1J4 | Cell shape-determining protein MreC | 29.7 | Regulation of Cell Shape [0008360] |
| A0A098Z983 | Phospho-N-acetylmuramoyl-pentapeptide-transferase | 36 | Cell Division [0051301], Cell Wall Organization [0071555], Peptidoglycan Biosynthetic Process [0009252], Regulation of Cell Shape [0008360] |
| A0A0B7LLG7 | Undecaprenyl-diphosphatase | 31.7 | Cell Wall Organization [0071555], Peptidoglycan Biosynthetic Process [0009252], Regulation of Cell Shape [0008360] |
| A0A0H2ZM82 | Cell division ATP-binding protein FtsE | 25.8 | Cell Division [0051301] |
| A0A0I7YM98 | Cell cycle protein GpsB | 13.1 | Cell Division [0051301], Regulation of Cell Shape [0008360] |
| A0A4F3K9I6, B8ZLT0 | Probable GTP-binding protein EngB | 22.2 | Division Septum Assembly [0000917] |
| A0A822Q745, A0A064C6I1 | Cell division protein DivIVA | 33.2 | Cell Division [0051301] |
| A0A4J1QNE6 | UDP-N-acetylenolpyruvoylglucosamine reductase | 32.9 | Cell Division [0051301], Cell Wall Organization [0071555], Peptidoglycan Biosynthetic Process [0009252], Regulation of Cell Shape [0008360] |
| Q4JZ00,<br>Q4K2E2,<br>Q4K046 | Capsular polysaccharide<br>biosynthesis protein CpsC | 25.6 | Capsular Polysaccharide<br>Biosynthesis [0045227] |
| Q4JZ01 | Tyrosine-protein phosphatase | 28 | Capsular Polysaccharide<br>Biosynthesis [0045227] |
| Q4K2E2 | Capsular polysaccharide<br>biosynthesis protein CpsC | 25.7 | Capsular Polysaccharide<br>Biosynthesis [0045227] |
| Q97QE5 | Ribitol-5-phosphate<br>cytidyltransferase | 26.2 | Cell Wall Organization<br>[0071555] |
| Q9ZII6 | Tyrosine-protein kinase<br>CpsD | 24.8 | Capsular Polysaccharide<br>Biosynthesis [0045227] |
| A0A064C372 | Probable manganese-<br>dependent inorganic<br>pyrophosphatase | 33.5 |  |
| P66708 | DNA-directed RNA<br>polymerase subunit alpha | 34.2 |  |

### Sorafenib counters bacterial fitness leading to complement deposition and serum killing

Having established that sorafenib impairs bacterial growth through inhibition of StkP, we next examined its downstream effects on key virulence-associated phenotypes, including bacterial cellular invasion, membrane integrity, morphological alterations, and susceptibility to complement-mediated killing. We used the GFP-expressing *S. pneumoniae* TIGR4 reporter strain (T4-GFP) to assess bacterial invasion into human A549 lung epithelial cells. A549 cells were infected with T4-GFP pre-treated with 10 µM sorafenib for 3 h, and the proportion of GFP-positive host cells were quantified by flow cytometry. The selected concentration of sorafenib reflects the clinically achievable plasma levels in patients undergoing cancer therapy (Fucile *et al*, 2015). While infection with untreated or DMSO-treated bacteria resulted in a positive peak shift in the GFP signal, indicative of successful invasion, sorafenib-treated bacteria failed to invade, showing fluorescence levels comparable to uninfected controls (**Fig. 4A)** with a significant reduction in the percentage of infected cells (**Fig. 4B**). To rule out host cytotoxicity at the bactericidal concentrations, we performed a PrestoBlue cell viability assay on A549 cells treated with sorafenib or DMSO across a range of concentrations (2.5–10 µM). No significant cytotoxicity was observed (**Fig. S7A**). Cells treated with 5 µM staurosporine, a known inducer of apoptosis, showed a ∼50% reduction in viability and served as the positive control for cytotoxicity. Next, we examined whether sorafenib affects bacterial membrane permeability. Differential viability staining using SYTO9 (green), which stains all bacteria, and propidium iodide (PI, red), which penetrates only cells with compromised membranes, revealed increased double-positive stained population upon sorafenib treatment (**Fig. 4C**). In contrast, untreated and DMSO-treated controls showed negligible PI staining. These results suggest that sorafenib induces significant membrane damage, which was further confirmed by flow cytometric analysis showing a significantly higher proportion of PI-positive cells in the sorafenib-treated group (**Fig. S7B**). To visualize the effects of sorafenib on bacterial morphology, we performed scanning electron microscopy analysis of bacteria grown in the presence of 10 μM sorafenib. Bacteria treated with the solvent alone, DMSO had intact cell wall (**Fig. 4D**), while sorafenib treated bacteria exhibited abnormal morphology with surface deformations indicative of compromised cell wall **(Fig. 4E).** Given the observed membrane disruption, we next assessed the susceptibility of sorafenib treated bacteria to complement-mediated opsonization. Bacteria treated with sorafenib, DMSO, or left untreated were incubated with human serum from healthy donors and complement protein C3 bound to bacterial membrane was quantified by flow cytometry. Sorafenib-treated bacteria showed significantly increased C3 deposition when compared to DMSO or untreated groups (**Fig. 4F, G**) both of which exhibited minimal complement binding due to the protective polysaccharide capsule. Notably, the level of C3 deposition on sorafenib-treated bacteria was comparable to capsule-permeabilized bacteria that were opsonized with anti-*S. pneumoniae* antibody, that was used as the positive control. Bacteria incubated with heat-inactivated serum served as the negative control and showed negligible C3 binding.

**FIGURE 4.**
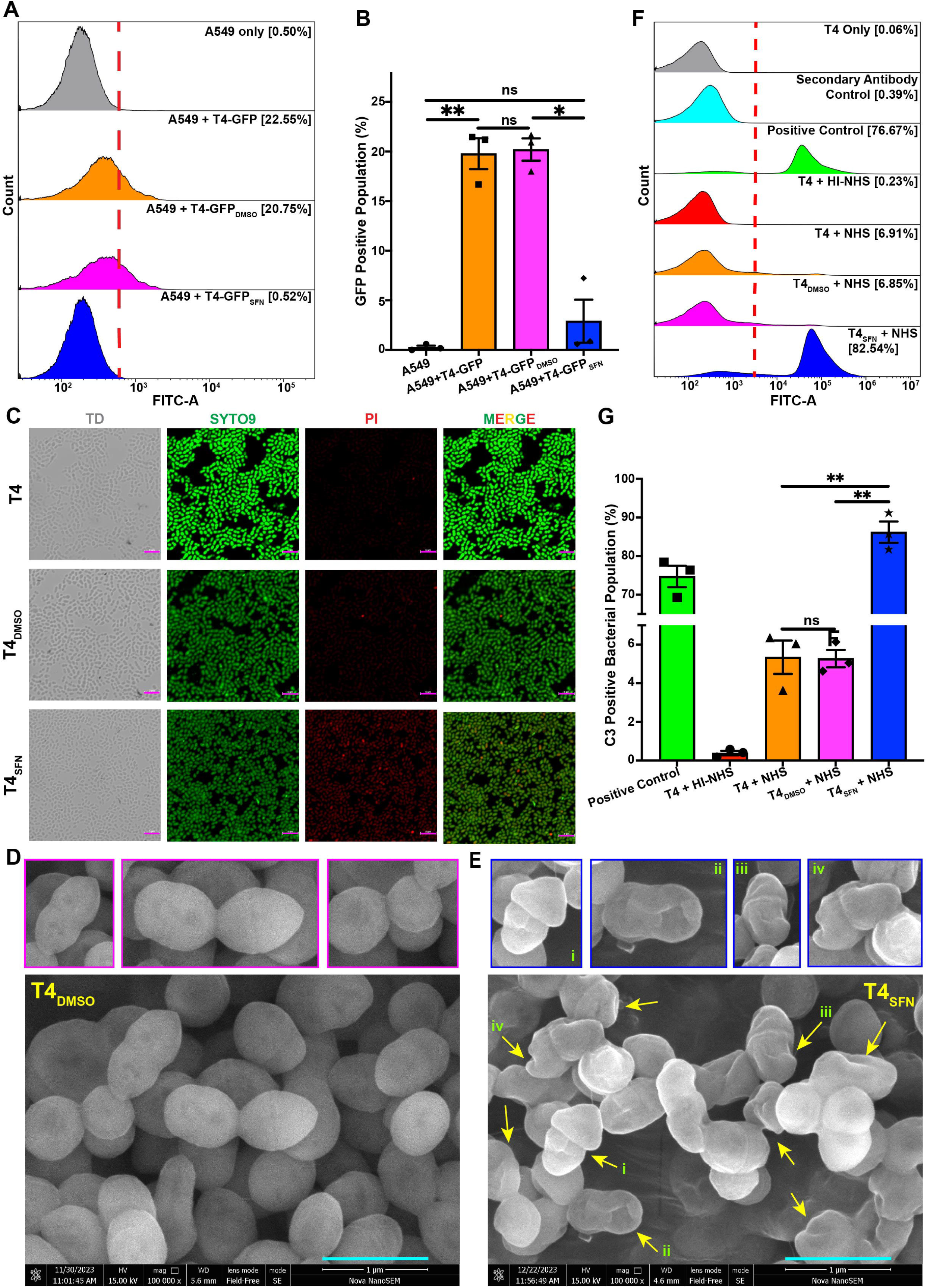
Sorafenib counters bacterial fitness leading to complement deposition and serum killing. **(A).** Flow cytometry histograms of A549 pneumocytes infected with GFP-expressing *S. pneumoniae* TIGR4 strain-(T4-GFP) strain grown in the presence or absence of 10 μM sorafenib (SFN). Treatment with equimolar concentrations of DMSO served as control. **(B)** Quantification of percentage of GFP positive infected cells in panel A showing reduced infection of A549 cells by SFN treated bacteria. * indicates p ≤ 0.05; ** indicates p ≤ 0.01; ns denotes non-significance by Welch’s t-test. **(C).** Viability staining of TIGR4 strain treated with 10 μM SFN using SYTO9 and PI dyes showing differential staining of live (green) and dead (red and green) bacteria. Scale bars, 5 µm. **(D, E).** Scanning electron microscopy of (**D**) 10 μM DMSO (solvent) and (**E**) SFN treated bacteria showing areas of compromised cell wall. Inset shows magnification of regions selected. Magnification 100,000x. Scale bars, 1 µm. **(F).** Flow cytometry histogram and (**G**) quantification analysis of complement C3 protein deposition on 10 μM SFN treated T4 bacteria upon incubation with normal healthy serum (NHS). Heat inactivated serum (HI-NHS) was used as negative control. Capsule permeabilized bacteria that were opsonized with anti-*Streptococcus pneumoniae* antibody was used as the positive control. ** indicates p ≤ 0.01 and ns denotes non-significance by Paired normal t-test. All data and images are representative of mean ± SEM from three independent experiments.

To test whether the enhanced complement deposition observed upon sorafenib treatment translated into higher bacterial killing, bacterial CFUs were determined upon incubation with human serum. As expected, untreated and DMSO treated bacteria incubated with serum, showed similar CFU counts regardless of serum exposure, indicating negligible serum-mediated killing under these conditions. In contrast, sorafenib-treated bacteria showed a significant reduction in CFU counts both in the presence and absence of serum, consistent with direct bactericidal activity of sorafenib (**Fig. S7C**). Collectively, these findings demonstrate that sorafenib not only inhibits StkP-activity but also compromises key bacterial virulence traits, including host cell invasion and immune evasion, by inducing membrane disruption and morphological defects, ultimately rendering *S. pneumoniae* highly susceptible to complement-mediated opsonization and killing.

### Limited Potential for Resistance Emergence upon Serial Passaging with Sorafenib

To evaluate the potential for resistance development against sorafenib, we performed serial passaging of *S. pneumoniae* in the presence of the drug. Bacteria were sequentially cultured on agar plates containing either the MIC concentration (2.5 µM) and sub-MIC concentration (1 µM) of sorafenib, and CFUs were quantified at each passage using plating assays (**Fig. 5A**). Following three consecutive passages at 2.5 µM, bacterial CFUs declined below the limit of detection, indicating an inability to propagate under continued drug pressure (**Fig. 5B**). Due to growth limitation at the MIC concentration, extended serial passaging (up to 16 cycles) was carried out at the sub-MIC level (1 µM). Throughout the experiment, no change in the MIC value was observed (**Fig. 5C**). Furthermore, analysis of CFU recovery at each passage revealed that the output CFUs remained consistently low, never exceeding 10-15% of the input CFUs by the end of passage 16 with no change in the MIC value (**Fig. 5D**). Collectively, these results suggest that prolonged exposure to sub-inhibitory concentrations of sorafenib does not promote resistance development in *S. pneumoniae* under the tested conditions.

**FIGURE 5:**
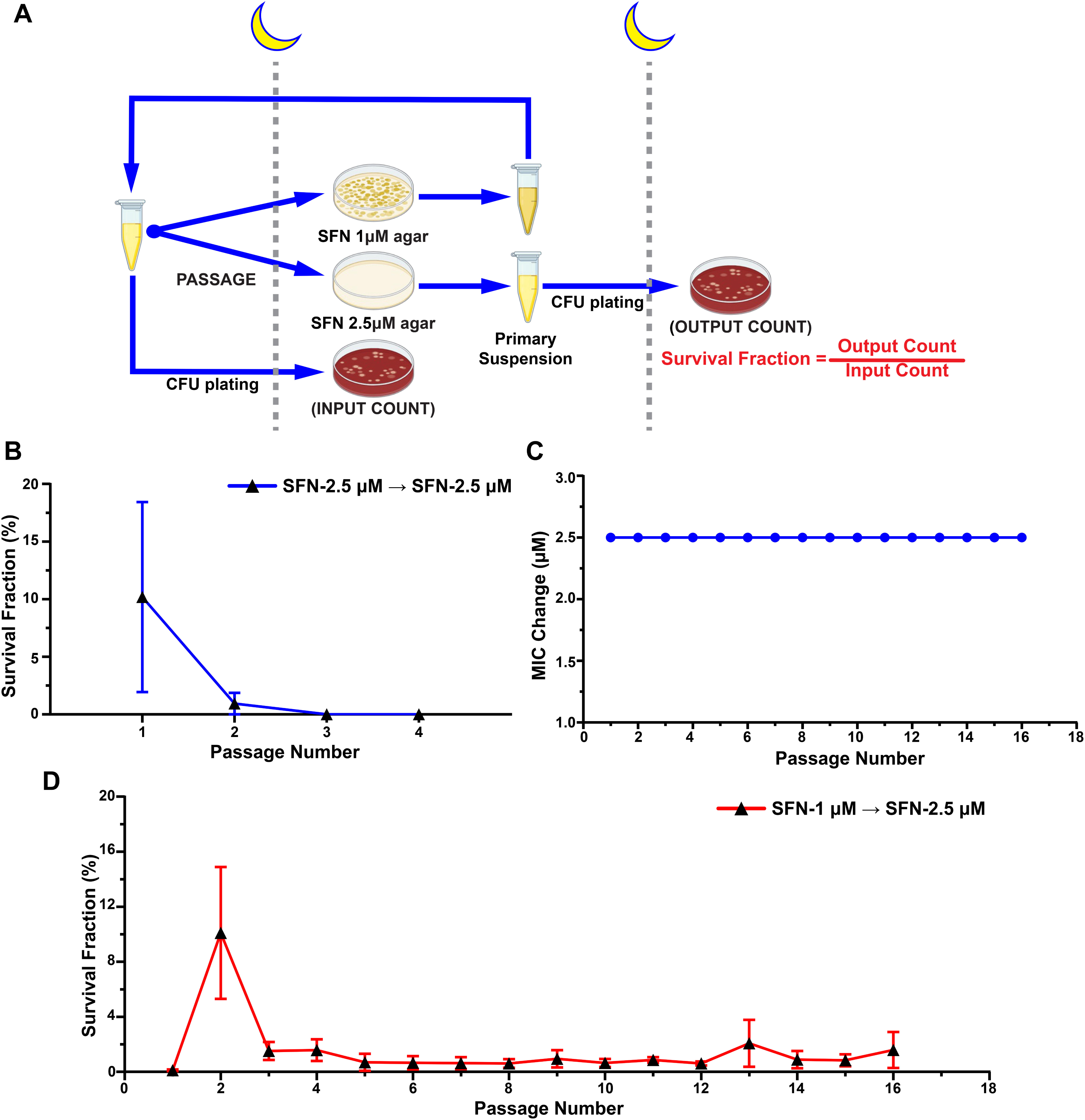
Limited potential for resistance emergence upon serial passaging with sorafenib. **(A).** Schematic showing the strategy of serial passaging of T4 strain with sub-MIC (1 µM) and MIC (2.5 µM) doses of sorafenib (SFN), followed by CFU enumeration of the respective plates. The passaging was performed 4x under 2.5 µM dose after which no CFU was recovered and 16x in 1 µM dose **(B).** Bacterial survival fraction at each passage upon serial passaging of T4 on BHI agar plates containing 2.5 μM (MIC) of SFN showing gradual decline up to passage 3 after which no CFU was recovered. The ratio of output CFU recovered at end of passage to the input from the previous passage was used to calculate the survival fraction. **(C)** Change in MIC upon serial passaging at 1 μM SFN and plating in BHI agar supplemented with 2.5 μM SFN. **(D).** Bacterial survival fraction showing viable pneumococci recovered on 2.5 μM SFN plates post successive passages with 1 μM SFN. All data are representative of mean ± SEM from three independent experiments.

### Sorafenib treatment reduces mortality and bacterial load *in vivo*

We next evaluated the *in vivo* efficacy of sorafenib in a murine model of pneumococcal pneumonia. Pneumonia was induced in 6–9-week-old male C57BL/6 mice via oropharyngeal administration of 1×10⁶ CFU of the *S. pneumoniae* serotype 4 strain, TIGR4. Mice in the treatment group received an initial intravenous dose of sorafenib (10 mg/kg in 50 µL of 55% PEG-400 and 20% DMSO) at 1 h post-infection, followed by intraperitoneal doses at 24 h intervals until the ethical endpoint was reached (**Fig. 6A**). The 10 mg/kg dose was selected to mimic a clinically relevant low-dose sorafenib regimen (equivalent to ∼45 mg/kg in human), which represents ∼10% of the dose typically used for hepatocellular carcinoma treatment (Reagan-Shaw *et al*, 2008) (Reiss *et al*, 2017), and is approximately one-third of the reported maximum tolerated dose in mice (Zaafar *et al*, 2024). Placebo mice received an equivalent volume of the vehicle (55% PEG-400 and 20% DMSO). Sorafenib-treated mice exhibited a ∼30% increase in survival compared to placebo-treated controls by day 4 post-infection, whereas all mice in the placebo group succumbed by day 3 (**Fig. 6B**). Consistent with enhanced survival, lung homogenates from sorafenib-treated mice showed ∼10-fold reduction in bacterial burden relative to control (**Fig. 6C**). Collectively, these results demonstrate that sorafenib treatment reduces pulmonary bacterial load and improves survival outcomes in a mouse model of pneumococcal pneumonia.

**FIGURE 6:**
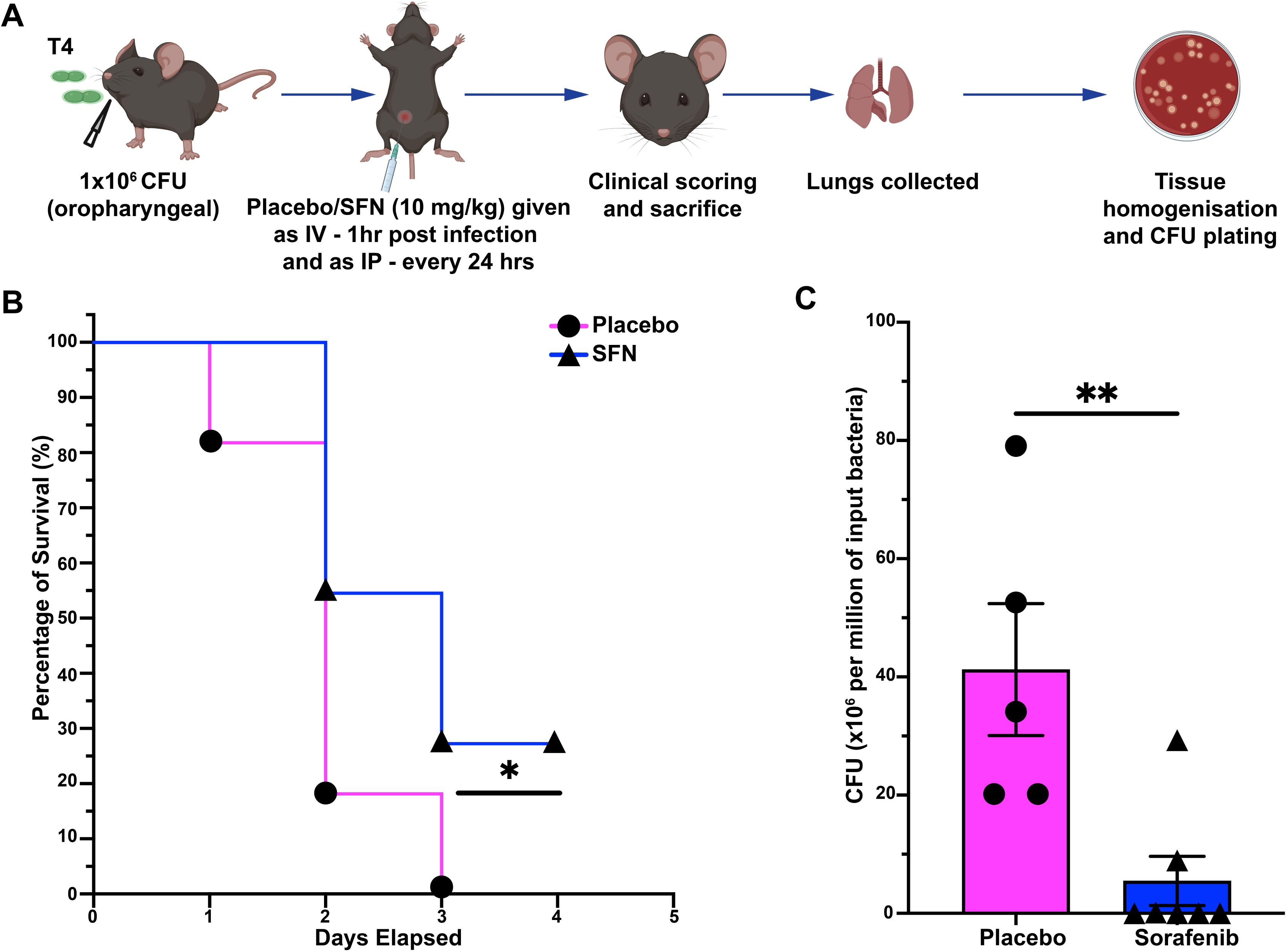
Sorafenib treatment reduces mortality and bacterial load *in vivo*. **(A).** Schematic showing the experimental workflow for testing the efficacy of sorafenib in six-to nine-week-old male C57BL/6 mice infected with 1×10^6^ CFU of TIGR4 strain via oropharyngeal route, followed by treatment with sorafenib (SFN) (10 mg/kg) intravenously at 1 h post infection and intraperitoneally at 24 h intervals. Infected mice treated with the solvent, 55% PEG-400 and 20% DMSO was used as the placebo control. **(B).** Survival curve of C57BL/6 mice (n=11) infected with T4 strain and treated with SFN or placebo over 4 days post infection. Infected mice were scored for clinical symptoms daily and were sacrificed upon reaching the ethical end point. * indicates p ≤ 0.05 by Mantel-Cox test. **(C).** Bacterial CFU (in million) recovered from the lungs of infected mice (*n* =5 for placebo; n=7 for SFN) was measured post-sacrifice and expressed normalized per million of input bacteria. ** indicates p ≤ 0.01 by Mann-Whitney test. Data are representative of mean ± SEM.

## Discussion

The growing threat of antimicrobial resistance in *Streptococcus pneumoniae* underscores the urgent need for novel therapeutic strategies. In this context, targeting conserved bacterial virulence factors, such as serine/threonine kinases offers a promising alternative strategy. Our study demonstrates the repurposing potential of sorafenib, a clinically approved multi-kinase inhibitor currently used in the treatment of thyroid, renal, and hepatocellular carcinomas (Raut *et al*, 2022) for treating pneumococcal infections and identifies the serine threonine kinase, StkP a potential target. StkP plays a multifaceted role in pneumococcal biology, including cell wall synthesis, stress response, competence, and cell division, making it an attractive target for virulence attenuation. The conservation of StkP among pneumococcal strains and the presence of homologous PASTA kinases (e.g., PknB in *Staphylococcus aureus* and *Mycobacterium tuberculosis*) further suggests potential for broader antimicrobial application. Our work builds on earlier studies targeting PASTA kinases in other bacterial pathogens, which found that selective inhibition of the *Staphylococcus aureus* kinase PknB (Tamber *et al*, 2010) and *Listeria monocytogenes* kinase PrkA (Pensinger *et al*, 2016) enhances susceptibility to beta-lactam antibiotics.

We identified sorafenib through *in silico* screening of host-directed kinase inhibitors against the StkP kinase domain. Molecular dynamics simulations revealed stable binding of sorafenib within the catalytic cleft of StkP, competing with its natural substrate, DivIVA. These findings provided a mechanistic basis for StkP inhibition by sorafenib. Crucially, we also demonstrated functional inhibition of StkP kinase activity through *in vitro* biochemical assays. Additionally, global phospho-threonine profiling of pneumococcal proteins following sorafenib treatment revealed a broad decrease in phosphorylation levels, consistent with StkP inhibition. The proteins with significant downregulation of phosphorylation were identified by mass-spectrometry. Functional validation demonstrated that sorafenib exhibited dose-dependent antimicrobial activity against both laboratory and clinical pneumococcal strains. In two highly erythromycin-resistant strains (GLO00007 and SP676), sorafenib displayed a ∼4-fold lower MIC compared to erythromycin, highlighting its effectiveness against antibiotic-resistant strains. Importantly, sorafenib treatment impaired bacterial adherence and invasion of human alveolar epithelial cells, consistent with StkP’s known role in regulating pneumococcal virulence (Echenique *et al*., 2004; Kant *et al*, 2023). Additionally, treated bacteria exhibited morphological abnormalities and compromised membrane integrity, phenocopying StkP effector mutants such as DivIVA, and GpsB (Fleurie *et al*., 2014). These defects were associated with increased C3 complement deposition and enhanced serum-mediated killing, indicating that StkP inhibition can also render bacteria more susceptible to host immunity. The complement pathway is critical for defense against invasive pneumococcal infections (Brown *et al*, 2002; Woehrl *et al*, 2011), and its enhanced activation may further potentiate therapeutic benefit. Importantly, minimal cytotoxicity was observed at effective antibacterial concentrations of 2.5-10 µM. These concentrations are within the range of human plasma levels achieved during cancer therapy of ∼12 µM in pediatric patients with solid tumors and refractory leukemias, administered with 200 mg/m^2^ dose (Widemann *et al*, 2012) and up to 20 µM in adults receiving a 400 mg dose (Mammatas *et al*, 2020) suggesting translational feasibility.

The specificity of sorafenib’s action on StkP was confirmed through genetic rescue assays. Ectopic expression of StkP in both wild-type and knockout backgrounds restored bacterial growth in the presence of sorafenib, establishing StkP as the target in *S. pneumoniae*. A critical concern in antimicrobial development is the potential for rapid resistance development. Serial passaging at sub-MIC concentrations up to 16 passages did not lead to an increase in MIC or bacterial regrowth, suggesting a low potential for rapid resistance development, an essential criterion for antimicrobial drugs. By targeting a conserved PASTA kinase, sorafenib introduces a novel mechanism distinct from traditional bactericidal or bacteriostatic drugs. This strategy disrupts regulatory signaling rather than essential metabolism or synthesis pathways, potentially imposing a higher fitness cost for resistance and reducing selective pressure, a concept gaining traction in anti-virulence drug development (Dickey *et al*, 2017) (Allen *et al*, 2014). *In vivo*, sorafenib demonstrated good efficacy in a murine model of pneumococcal pneumonia, improving survival and reducing lung bacterial burden at a low dose (10 mg/kg), which corresponds to approximately 10% of the clinical oncology dose and one-third of the maximal tolerated dose in mice. This is particularly important as invasive pneumococcal disease is associated with high morbidity and mortality rates. However, further studies with larger sample sizes and extended treatment durations are needed to fully elucidate the long-term effects and optimal dosing regimens of sorafenib in pneumococcal infections. Our study concurs with recent studies that reported sorafenib’s antimicrobial activity against MRSA (Chang *et al*, 2016; Le *et al*., 2020), but in these studies the role of bacterial kinase target was not identified.

In conclusion, this study provides compelling evidence that sorafenib, through inhibition of the conserved bacterial kinase StkP, impairs key virulence and survival mechanisms in *S. pneumoniae* **(Fig. 7)**. Targeting bacterial Ser/Thr kinases may represent a promising anti-virulence strategy to combat antimicrobial resistance. Further development and optimization of sorafenib and related kinase inhibitors targeting StkP could yield a new class of antimicrobials against pneumococci and other high-priority pathogens.

**FIGURE 7:**
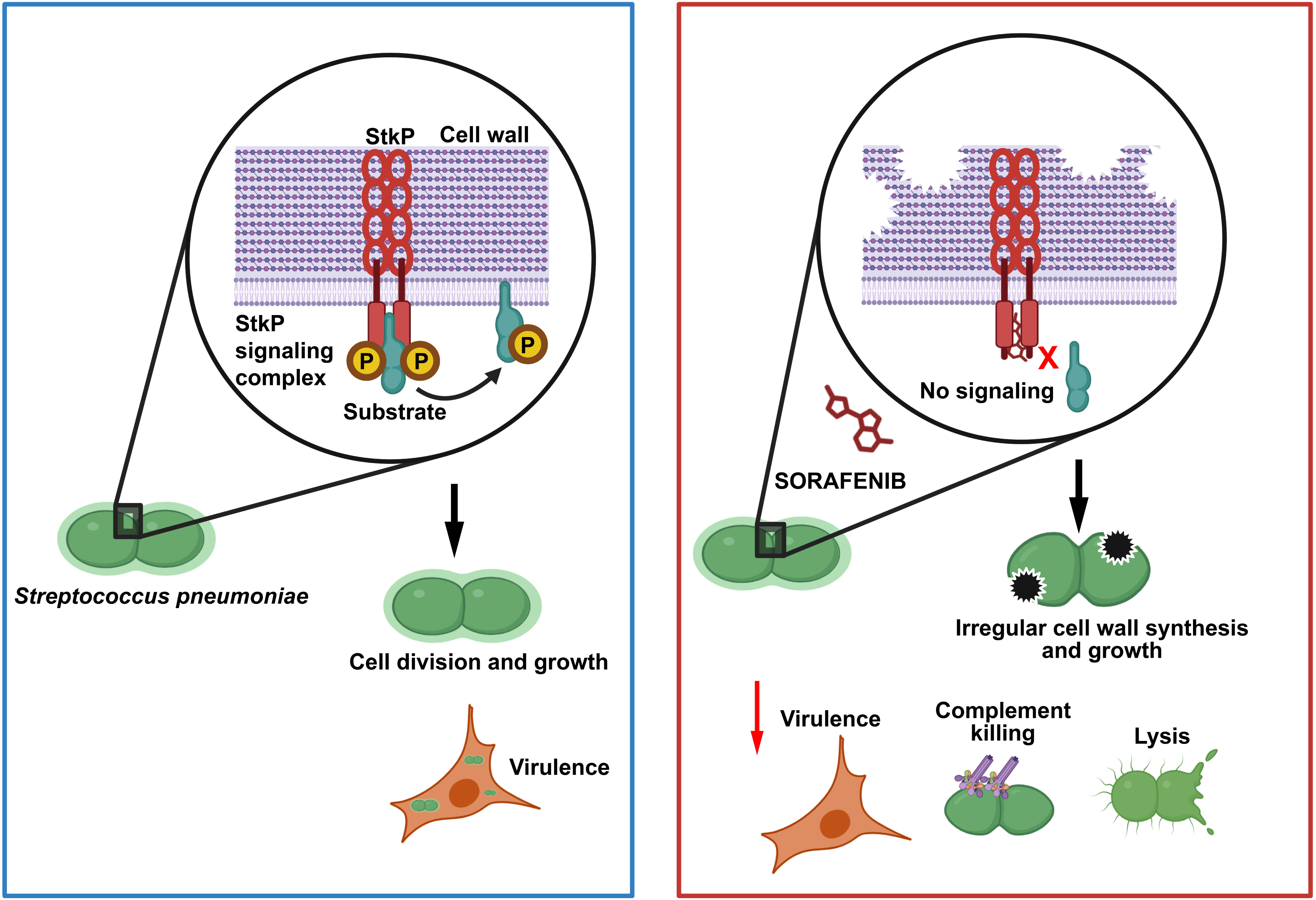
Sorafenib counters pneumococcal growth and virulence by targeting StkP. Schematic summarizing the antimicrobial effects of sorafenib against *Streptococcus pneumoniae*. The serine threonine kinase, StkP plays vital role in bacterial growth by coordinating cell wall synthesis and septal division. Our study suggests that sorafenib binds to the catalytic cleft of the StkP kinase domain, thereby inhibiting its kinase activity and resulting in irregular cell wall and enhanced membrane permeability. Blockade of StkP by sorafenib also reduced bacterial invasion into lung epithelial cells and enhanced susceptibility to complement mediated killing in serum. The *in vivo* efficacy was also validated in a mouse model of pneumonia.

## Acknowledgements

The study was supported by extramural funding through the INSPIRE Faculty fellowship Department of Science and Technology (DST/INSPIRE/04/2019/002238) and Ramalingaswami Re-entry fellowship (BT/RLF/Re-entry/46/2020) from the Department of Biotechnology, India and intramural funding from BRIC-Rajiv Gandhi Centre for Biotechnology, Thiruvananthapuram. We profusely thank Prof. Birgitta Henriques Normark, Karolinska Institutet, Stockholm and Prof. Anirban Banerjee, Indian Institute of Technology, Bombay for sharing pneumococcal strains. We also greatly acknowledge Dr. Pavel Branny, Czech Academy of Sciences, Czech Republic for gifting the StkP plasmids. We thank Dr. Kamalakannan Vijayan, School of Biology, Indian Institute of Science Education and Research, Thiruvananthapuram for sharing sorafenib and dasatinib for preliminary screening experiments.

We acknowledge Sonali Ghosal and Dr Dileep Vasudevan (Structural Biology Laboratory, BRIC-Rajiv Gandhi Centre for Biotechnology, Thiruvananthapuram) for their generous assistance with the StkP-KD protein purification.

## Author contributions

K.S. conceptualized the project. K.S, S.G. and B.V. acquired resources and supervised the study. J.A, A.C.S, H.D, C.P, S.A, P.M.K, A.R, K.B., A.N, R.V. and S.K.C. conducted the investigations. J.A, A.C.S, H.D, C.P, S.A, P.M.K, A.R, K.B., A.N, R.V, A.N, S.K.C, K.S and B.V. and analyzed the data. K.S. and J.A. wrote the original draft. K.S, J.A, S.G, B.V. and reviewed and edited the manuscript. K.S, S.G. and B.V. acquired funding. All authors have reviewed and approved the manuscript.

## Materials and Methods

### Protein structure prediction and assessment

The protein sequence data of *Streptococcus pneumoniae* (strain ATCC BAA-334 / TIGR4) StkP was obtained from the UNIPROT database (UniProt, 2025) under the accession code Q97PA9. Due to the nonavailability of crystal structure for the pneumococcal StkP, particularly its kinase domain spanning residues 12 to 273, we employed AlphaFold (Tunyasuvunakool *et al*, 2021) to generate a structural model for this domain. Specifically, we trimmed the structure for the residues 1 to 299 (StkP-KD), which was further processed using Schrödinger. Finally, we utilized the structure assessment tool within the SWISS-MODEL server (Waterhouse *et al*, 2024) to generate and analyze the Ramachandran plot of the constructed model that helps to evaluate the stereochemical quality and stability of the protein model.

Furthermore, we have used PyMOL (PyMOL Molecular Graphics System, Version 3.0 Schrödinger, LLC) for structural superimposition and visualization of the alignments and to calculate the Root Mean Square Deviation (RMSD) between the constructed model of StkP-KD and the corresponding structures of *Staphylococcus aureus* serine/threonine kinase protein – PknB (PDB ID: 4EQM) and *Mycobacterium tuberculosis* serine/threonine kinase proteins, PknB (PDB ID: 3ORI) and PknA (PDB ID: 4OW8) to quantify structural similarities and deviations. Sequence alignment diagram was made using the ESPript 3.0 (ENDscript - https://endscript.ibcp.fr) (Robert & Gouet, 2014).

### Virtual screening of ligands and docking with StkP

The prepared ligands were docked into the protein at the generated grid site using the extra precision (XP) docking algorithm in the Glide module (Friesner *et al*., 2006) [Schrödinger Release 2024-4: Glide, Schrödinger, LLC, New York, NY, 2024] of Schrödinger. Ligands were sampled flexibly, with nitrogen inversions and ring conformations considered, while macrocycles were not sampled, and the input ring conformation was excluded. Torsion sampling was biased for all predefined functional groups, and Epik state penalties were added to the docking score. No core, shape, or torsional constraints were applied during docking. Post-docking minimization was performed, limiting the number of poses per ligand to five. The output was saved as a pose viewer file, including the protein structure.

### Molecular dynamics simulations

The docked structure of the StkP-KD and sorafenib complex was loaded into Maestro and prepared using the Protein Preparation Wizard, as described previously for docking preparations. Molecular dynamics (MD) simulations were conducted using the Schrödinger Desmond software v.2023-2 (Bowers *et al*, 2006) [Schrödinger Release 2024-4; Desmond Molecular Dynamics System, New York, NY, 2024] for a duration of 200 ns, with frames recorded every 100 ps. An orthorhombic box was created around the prepared StkP-KD-sorafenib complex with a 10 Å buffer of solvent molecules using the System Builder module. The box volume was minimized, and ions were added to neutralize the system. TIP3P solvent molecules were used for solvation (Jorgensen *et al*, 1983). The system’s temperature was maintained at 300K using the Nose-Hoover chain thermostat (Hoover, 1985) with a relaxation time of 1.0 ps, while the pressure was kept at 1.01 bar using the Martyna-Tobias-Klein barostat (Martyna *et al*, 1994) with isotropic coupling and a relaxation time of 2.0 ps. Coulombic interactions were handled using a short-range cutoff method with a cutoff distance of 9.0 Å. The OPLS4 force field was used to describe the interactions within the system.

DivIVA is a widely phosphorylated natural substrate of pneumococcal StkP (Novakova *et al*., 2010). The full-length protein is 262 amino acids and Thr-201 is an important residue that is phosphorylated by StkP (UNIPROT ID: Q8CWP9) (Fleurie *et al*., 2012). Since the crystal structure of DivIVA is not available, we processed the AlphaFold predicted structure of DivIVA to extract a nine amino acid containing peptide centered at Thr-201. The peptide was added near the catalytic cleft of StkP-KD complexed with sorafenib and prepared using the Protein Preparation Wizard. The interactions were simulated for 200 ns as previously described.

### Prime – MM/GBSA calculations

The binding affinity of the ligand from MD simulation trajectories was calculated using the Molecular Mechanics-Generalized Born Surface Area (MM/GBSA) method implemented in the Prime module of Schrödinger Suite (Jacobson *et al*, 2004). The out.cms file from the MD simulation, which contains the trajectory data, was used as the input along with the specified ligand residue number. The binding free energy, which measures the thermodynamic stability of the protein-ligand interaction was calculated using the below equation:

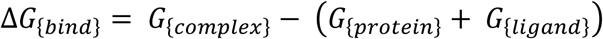

Here, G_complex_ is the free energy of the protein-ligand complex, G_protein_ is the free energy of the unbound protein, and G_ligand_ is the free energy of the unbound ligand.

### Bacterial strains and culture

The encapsulated strains of *S. pneumoniae* serotype 4, TIGR4 (T4; ATCC BAA-334) and serotype 2, D39 (NCTC 7466) were kindly gifted by Prof. Birgitta Henriques Normark, Karolinska Institutet, Stockholm. The GFP expressing T4 strain, T4-GFP, was kind gift from Prof. Anirban Banerjee, Indian Institute of Technology, Bombay. The D39-gfp-stkP(P_czcD_ = P_Zn_) strain containing an additional inducible copy of stkP gene inserted in the genome at the dispensable beta-galactosidase (bgaA) locus and the D39ΔstkP::gfp-stkP(P_czcD_ = P_Zn_) strain with endogenous stkP gene knocked out were made in the lab and used for the experiments.

The TIGR4 and D39 strains were streaked from frozen glycerol stocks, on soyabean casein digest agar plates supplemented with 5% sheep blood (HiMedia) and grown at 37°C and 5% CO_2_ overnight. T4-GFP strain was grown in BHI agar (HiMedia) supplemented with 4 μg/mL of chloramphenicol (Sigma-Aldrich). D39-gfp-stkP(P_Zn_) was streaked on BHI agar plate containing 2.5 μg/mL of tetracycline (HiMedia). To induce the gfp-stkP gene under the P_czcD_ promoter, the strain was streaked on BHI agar plate containing 2.5 μg/mL of tetracycline, 0.5 mM of Zn^2+^ (as ZnCl_2_) (HiMedia), and 0.05 mM of Mn^2+^ (as Mn(II)Cl_2_) (HiMedia). The D39ΔstkP::gfp-stkP(P_Zn_) strain was streaked on BHI agar plates containing 2.5 μg/mL of tetracycline, 4.5 μg/mL of chloramphenicol, 0.5 mM of Zn^2+^ and 0.05 mM of Mn^2+^. All strains were grown in Brain-Heart Infusion (BHI) broth (HiMedia) for suspension cultures with appropriate antibiotics and inducing reagents.

For all inhibitor-based assays, Sorafenib (Sigma-Aldrich) was dissolved in 100% DMSO (HiMedia) to make 100 mM stocks. This stock was further diluted to 10 μM in BHI media and used, effectively making the final DMSO concentration to 0.01%, less than the known cytotoxic levels. Also, equimolar concentrations of DMSO were used as the solvent control. All experiments were performed with prior approval from the Institutional biosafety committee, BRIC-RGCB (Ref. 51/IBSC/karthik/202151) and Review Committee on Genetic Manipulation, Department of Biotechnology, Govt. of India (Ref. BT/IBKP/363/2020).

### Cell culture

Human adenocarcinoma alveolar basal epithelial cells, A549 (ATCC CCL-185), were procured from the National Centre for Cell Sciences, Pune, India. The cell lines were authenticated using a short tandem repeat 10 analysis and tested negative for mycoplasma contamination. A549 cells were grown in Dulbecco’s Modified Eagle Medium (DMEM) media (HiMedia) supplemented by 10% Fetal Bovine Serum (FBS) (GIBCO) and 1% penicillin-streptomycin antibiotic (HiMedia) and were incubated at 37°C and 5% CO_2_.

### Bacterial growth kinetics assay

To measure the effect of sorafenib on pneumococcal growth, a kinetic assay was set up to measure OD_620_ using the MultiSkan FC Microplate reader (ThermoFisher). Serial dilutions of both DMSO and sorafenib were made in BHI media and was mixed with equal volume of *S. pneumoniae,* TIGR4 and D39 culture to give a starting OD_620_ of 0.05-0.06. 100 μL of each was added as triplicates into a flat bottom 96-well plate (Tarsons) and inserted into the plate reader after sealing with an optically transparent sheet (Tarsons). The plate was incubated at 37°C within the instrument and OD_620_ was automatedly measured every 20 min interval for 18 h. Bacteria incubated in BHI media alone was used as untreated control.

### MIC determination in clinical strains

The minimum inhibitory concentration (MIC) of the sorafenib and the comparators (Penicillin, Erythromycin, and DMSO) were determined by broth microdilution method as recommended by Clinical Laboratory Standard Institute (CLSI) (M07, 2019). Quality control strain *S. pneumoniae* ATCC 49619 was included in each run. Six clinical strains with varying penicillin and erythromycin MIC values as listed in **Table 1** were used for MIC testing. Briefly, the bacterial inoculum (10^5^ CFU/mL) was prepared in Cation Adjusted Mueller Hinton Broth (CAMHB) with 2.5% Lysed Horse Blood (LHB). The stock sorafenib (100 mM =46.46 mg/mL) and the comparator antimicrobials were diluted in CAMHB with LHB to get 128 μg/mL concentration and then serially diluted in the microtiter plate up to 0.06 μg/mL. Equal volume of the inoculum (50 μL) was added to each well containing the serially diluted drugs in a final volume of 100 μL. The bacterial concentration of the inoculum that was added into the titer plate was calculated by performing serial dilution from the growth control. The plates were incubated at 37°C, for 20-24 h. The wells were tested the next day for the inhibition of growth and the viability (bactericidal effect). The MIC value of sorafenib /antibiotic is taken as the lowest concentration which shows the visible inhibition of the growth. The susceptibilities of antibiotics were interpreted following the CLSI criteria (M100, 2019). The viability was measured in CFU/mL by subculturing from four concentrations (64, 32, 16 and 8 μg/mL); one dilution above and two dilutions below the MIC, and from the MIC well on to sorafenib/antibiotic free media the next day after incubation. To determine the CFU/mL, the serial dilutions were made from each concentration and inoculated onto blood agar plates for colony count.

### Generation and validation of gfp-stkP overexpressing and complemented mutant strains

*S. pneumoniae*, D39 strain was induced to competence by stimulation with 10 µg/mL of CSP-I (AnaSpec) in BHI competence media that comprised of 10 mL BHI broth, 100 μL of 100 mM CaCl_2_, and 250 μL of 8% BSA at pH 8.0 for 10 min at 30°C. The competent cells were transformed with 1 μg of pJWV25-stkP plasmid (kindly gifted by Dr. Pavel Branny, Czech Academy of Sciences, Czech Republic) to create D39-gfp-stkP(P_Zn_). The plasmid stably integrates into bacterial genome by homologous recombination at the dispensable bgaA gene under a zinc inducible promoter gene P_czcD_ (≈P_Zn_). The mixture was then incubated for 20 min at 30°C and then for 2 h at 37°C. After incubation, the cell suspension was spun down at 5000 rpm for 5 min and plated on BHI agar containing tetracycline at 2.5 μg/mL concentration. D39-WT without any plasmid DNA was used as a control. The positive clones were grown in BHI media supplemented with 2.5 μg/mL tetracycline to OD_600_ of 0.1. The gfp-stkP gene was induced by adding 0.5 mM Zn^2+^ (as ZnCl_2_) and 0.05 mM Mn^2+^ (as Mn(II)Cl_2_) as reported previously (Beilharz *et al*., 2012; Eberhardt *et al*, 2009). The cultures were grown to mid-log phase and was pelleted down. The GFP-StkP protein expression was validated using the CytoFlex S Flow Cytometer (Beckman-Coulter). The data were analyzed using Kaluza version 2.2.1. The protein localization was validated by using a Nikon Eclipse Ti2 Inverted Confocal Imaging System. D39-WT and D39-gfp-stkP(P_Zn_) grown in the absence of Zn^2+^ and Mn^2+^ were used as negative controls.

The protein expression was also confirmed by western blotting. D39-WT and D39-gfp-stkP(P_Zn_) pellets were lysed using RIPA lysis buffer (Himedia) containing Protease Inhibitor Cocktail (Roche) on ice for 30 min. 30 μL of the crude lysate was mixed with 10 μL of 4X Lamelli Buffer containing 50 mM DTT, heated for 10 min at 70°C and loaded on a 12% SDS-PAGE. The proteins were transferred to a PVDF membrane using a Transblot Turbo Transfer System (Bio-Rad) at 1.3 A and 25 V for 15 min. The membrane was blocked using 5% BSA (HiMedia) in TBS buffer for 1 h at room temperature and then incubated with goat-induced anti-GFP antibody (Santa Cruz) at 1:400 dilution for overnight at 4°C. The membrane was collected after the incubation and was washed twice for 5 min and thrice for 10 min with TBS buffer containing 0.1% Tween-20 (Bio-Rad) (0.1% TBST). The membrane was then incubated with HRP-conjugated anti-goat secondary antibody (Bio-Rad) at 1:2500 dilution and StrepTactin-HRP Conjugated secondary (Bio-Rad) at 1:10,000 dilution to detect *Strep*-tagged Precision Plus Protein™ WesternC Blotting Standards, Bio-Rad for 1 h at room temperature. The membrane was washed twice for 5 min and twice for 15 min with 0.1% TBST, then subjected to a final 15 min wash in TBS buffer containing 0.5% Tween-20. The membrane was developed using Clarity Western ECL Substrates (Bio-Rad) and the luminescence was acquired using iBright FL1500 Imaging System (Invitrogen).

To create the complemented mutant strain, D39ΔstkP::gfp-stkP(P_Zn_), the D39-gfp-stkP(P_Zn_) strain was transformed with 1 μg of AkCmDkS plasmid (gifted by Dr. Pavel Branny, Czech Academy of Sciences, Czech Republic), which replaces the endogenous stkP gene with chloramphenicol resistant gene by homologous recombination. The colonies were plated on BHI agar containing 2.5 μg/mL of tetracycline, 4.5 μg/mL of chloramphenicol, 0.5 mM of Zn^2+^, and 0.05 mM of Mn^2+^. D39-gfp-stkP(P_Zn_) cells without any plasmid DNA was used as a control. The positive clones were verified by flow cytometry and confocal microscopy as described earlier. The stkP gene copies in D39-gfp-stkP(P_Zn_) and D39ΔstkP::gfp-StkP(P_Zn_) respectively were validated by PCR with 5’-CCTCTGCAACTGTCTGACC-3’and 5’-CAGATGGGACTGCCAAGG-3’ as the forward and reverse primers respectively. 250 ng of genomic DNA was used as the template, and the PCR reaction was performed for 13 cycles to eliminate the band saturation effects.

### Phospho-Threonine western blotting

Overnight grown colonies of D39 and T4 strains were made into a suspension and diluted to OD_600_ of 0.1 in BHI media. DMSO and sorafenib were added to the cultures at the required concentrations. The cultures were incubated at 37°C in a water bath. Post 4-5 h of treatment, cultures were spun at 10,000 rpm speed and 4°C. The pellets were washed once with ice cold PBS, followed by treatment with RIPA lysis buffer supplemented with Protease Inhibitor Cocktail and PhosSTOP (Roche) and were incubated on ice for 2-3 h for the lysis reaction. The lysate protein concentration was quantified by BCA Protein Assay Kit (Pierce). 15 μg of the lysate was resolved on a 12% SDS-PAGE. Rabbit induced anti-*S. pneumoniae* enolase antibody at 1:50,000 dilution in blocking buffer (Gifted by Prof. Anirban Banerjee, Bacterial Pathogenesis Laboratory, Indian Institute of Technology, Bombay, India) was used as the loading control. Mouse anti-Phospho-Threonine monoclonal antibody (CST) at 1:1000 dilution in blocking buffer supplemented with 0.1% Tween-20 was used for probing the phosphorylation pattern. Goat induced anti-rabbit and anti-mouse antibodies conjugated to HRP (Bio-Rad) were used as the secondary antibodies respectively.

### StkP phosphorylation in *E. coli*

*E. coli* BL21(DE3) cells transformed with KDNC2 plasmid construct were seeded into primary culture in Luria Bertani (LB) broth supplemented with 100 μg/mL of ampicillin and incubated at 30°C at 200 rpm for 16 h. For the secondary culture, 1% primary suspension was added to LB broth supplemented with 100 μg/mL of ampicillin. Sorafenib and equivalent amount of DMSO was added into the secondary culture at the specified concentrations and incubated at 37°C at 200 rpm for 4 h. The culture was spun at 10,000 rpm at 4°C and washed with ice cold PBS. The pellets were lysed using RIPA buffer supplemented with Protease Inhibitor Cocktail and PhosSTOP. The lysate was incubated in ice for 2-3 h with periodic gentle vortex. The lysates were then sonicated for 2 min time with 2 s on and 3 s off cycles at 30% amplitude, followed by centrifugation at 16,000g for 30 min at 4°C. The supernatant was collected and stored in 4°C for further analysis. Total protein of the supernatant was estimated by BCA Protein Assay Kit (Pierce) and 20 μg of the total protein was resolved on a 12% SDS-PAGE. Coomassie stained gel was used as the loading control. Western blotting with anti-Phospho-Threonine monoclonal antibody was performed as described earlier, but with washes using TBS buffer containing 0.5% Tween-20. BL21(DE3) cells transformed with empty pET-22b(+) vector and KDNC2 transformed cells supplemented with 0.5 mM IPTG at OD_600_=0.5 for StkP-KD induction were used as the negative and positive controls for StkP-KD respectively. Only 2 μg of the clarified lysate was loaded for StkP-KD induced sample to minimize band saturation upon StkP induction.

### A549 infection assay

*S. pneumoniae* TIGR4-GFP strain was grown on BHI agar plates containing chloramphenicol (CmR). Isolated colonies were inoculated in BHI broth containing CmR till OD_600_ reached 0.1. The cultures were treated with 10 µM Sorafenib and DMSO and incubated in the water bath at 37°C until OD_600_ reached 0.4. Based on the CFU count, i.e. 3.55 x 10^7^ CFU/mL, the volume of bacterial suspension containing 2 x 10^7^ cells/well for M.O.I of 100:1 was washed and resuspended in 50 µL of PBS. A549 cells were seeded in a 12-well plate (seeding density 2 x 10^5^ cells/well) in 750 µL Dulbecco’s Modified Eagle Medium (DMEM) (HiMedia) supplemented with 10% fetal bovine serum (FBS) (Invitrogen) and incubated overnight at 37°C, 5% CO_2_. Untreated, DMSO-treated, and sorafenib-treated bacteria were added to A549 cells containing wells with 750 µL DMEM supplemented with 5% heat-inactivated FBS without antibiotics. The plate was spun at 1500 rpm for 5 min to enhance bacterial contact and incubate at 37°C, 5% CO_2_ for 3 h. After incubation, the media was aspirated gently and A549 cells were harvested by trypsinization. The cells were washed and fixed at room temperature for 10 min with 4% paraformaldehyde. The cells were analyzed using the BD FACS Aria III cytometer (BD Biosciences) in the FITC channel to detect GFP-positive signals.

### Live-dead staining of bacteria

Live-Dead staining was performed using the BacLight Bacterial Viability Kit (ThermoFisher), following the manufacturer’s protocol. Bacteria were grown to mid-log phase in the presence of 10 μM Sorafenib and DMSO. Approximately 5.8 x 10^7^ CFU/mL of bacterial cells were washed and resuspended in 500 μL PBS. Each sample was incubated with 1.5 μL of the staining dye mix containing equal volumes of SYTO9 and Propidium Iodide for 15 min under dark conditions. The cells were washed twice in PBS again, followed by fixation with 4% paraformaldehyde (HiMedia) at room temperature for 10 min. Bacteria treated with 70% Ethanol (Molecular Biology Grade, Supelco) for 15 min at room temperature was used as the positive control of death and was spun down at 10,000 rpm for staining and further steps. The fixed cells were mounted (ProLong Glass Antifade Mountant, Invitrogen) on glass slides, using No. 1 English Coverslips (Bluestar) and were stored in 4°C till imaging. The slides were visualized using a Nikon Eclipse Ti2 Inverted Confocal Imaging System.

### Scanning electron microscopy

The bacteria were cultured in BHI broth at 37°C and 5% CO_2_ until an OD_620_ of 0.7 was reached. The bacteria were then pelleted and washed thrice with PBS, following fixation with 4% paraformaldehyde at room temperature for 2 h. After rinsing the pellet thrice in distilled water, sequential dehydration was followed using 30% ethanol for 10 min, 50% ethanol for 10 min, 70% ethanol for 10 min, and 100% ethanol for 15 min. Subsequently, a speed vacuum was used in 30-second intervals for a duration of 3 min, to obtain the bacterial pellet as a fine powder. This was then air-dried under a laminar flow for 1-2 h. The adhesive carbon tape (PELCO Tabs, 12mm OD, 16084-1, Ted Pella) was placed on the sample holders, and the powdered sample was dusted on the tape using a paintbrush. This was then sent for gold sputtering. The gold sputtering was done using a QT Quorum Sputter Coater with a thickness of about 10 nm. Gold-coated specimens were imaged, using FEI Nova Nanosem 450 at Indian Institutes of Science Education and Research, Thiruvananthapuram, under a high vacuum at 15.0 kV, with a 5.6 mm working distance and a 40 μm objective lens aperture. Images were collected using the secondary electron detector; the acquisition time per image was 10-20 μs, and each image was 1536 × 1103 pixels. SEM images were recorded at magnifications ranging from 10,000 x to 100,000x.

### Complement deposition and serum killing assay

To check the susceptibility of sorafenib treated pneumococci for the innate immune system, an assay was performed to quantify the deposition of complement C3b protein on the bacteria. Briefly, *S. pneumoniae* TIGR4 strain was grown to mid-log phase in BHI media. Untreated and 10 μM DMSO treated bacteria were grown to OD_600_=0.5 while the 10 μM sorafenib treated bacteria maintained a static OD_600_ of 0.1. Upon OD normalization of both cultures to 0.5 at OD_600_, ∼ 5.8*10^7^ CFU/mL of bacterial cells were washed with PBS buffer and and incubated with 10% serum (diluted in PBS) obtained from healthy human donors at 37 °C for 30 min. Pellets incubated with heat-inactivated human serum served as the negative control. Since TIGR4 strains are encapsulated, bacteria with capsule permeabilized by 0.05% Triton X-100 (HiMedia), followed by opsonization at 37°C for 30 min with anti-*Streptococcus pneumoniae* antibody (ab20429; Abcam) was used as the positive control. The bacterial-bound C3 protein was stained with goat induced anti-complement 3 primary antibody (Sigma-Aldrich, 204869) at 1:100 dilution, followed by an incubation with donkey induced anti-goat secondary antibody conjugated with Alexa Flour 488 (Invitrogen, A-11055) at 1:400 dilution. The bacteria were blocked with 1% BSA (HiMedia) in PBS before antibody incubation and all steps had one PBS wash in between. Finally, the samples were fixed with 2% paraformaldehyde (HiMedia) and resuspended in PBS. The samples were stored at 4°C till analyzing them by flow cytometry (CytoFlex S, Beckman-Coulter). The data were analyzed using Kaluza version 2.2.1.

For serum killing, the bacterial cells incubated with serum were washed and resuspended in 100 µL PBS, serially diluted and plated each dilution as three replicates on blood agar plates. The plates were incubated at 37°C and 5% CO_2_ and colonies were counted after 16 h.

### Serial passaging assay for drug resistance

Pneumococci were continuously passaged on plates containing 1 μM (sub-MIC) and 2.5 μM (MIC) concentrations of sorafenib to study resistance development. For sub-MIC passaging experiments, 50 μL of OD normalized pneumococci post pre-treatment with 1 μM sorafenib in BHI broth (Passage 0) were plated on BHI plates containing 1 μM and 2.5 μM of sorafenib (Passage 1). Post overnight incubation at 37°C and 5% CO_2_, the plates were assessed for growth. Colonies grown on SFN-1 μM plates were harvested into 2 mL of BHI broth to create a primary suspension. The primary suspension was OD-normalized to 0.5 and 50 µL was subsequently plated onto 1 μM and 2.5 μM sorafenib plates (Passage 2) and remaining suspension was frozen. For initial passages under 1 μM sorafenib showing minimal growth (and thus undetectable OD), one-fifth of the primary suspension was directly plated. To evaluate survival rates, OD-normalized pneumococci treated with 1 µM sorafenib were serially diluted and plated on blood agar. Colonies obtained from 50 µL of this dilution were counted and defined as the input CFU. For output CFU determination, 100 µL of the primary suspensions (adjusted to a final volume of 2 mL) collected from SFN-2.5 μM plates were serially diluted on blood agar. The procedure was carried out for 16 passages.

For MIC passaging experiments, one-fourth of the primary suspension obtained from SFN-2.5 μM plates were directly plated without OD normalization due to the negligible growth on the plates. Plating was continued for 4 passages until no colony growth was observed, indicating minimal resistance.

### Mouse experiments

Mouse experiments were approved by the institutional animal ethics committee (Ref. IAEC/924/KARTHIKS/2023). C57BL/6J mice were generated and maintained at the Animal Research Facility, BRIC-RGCB using Individually Ventilated Caging (IVC) system with 14 h light and 10 h darkness at 25°C with *ad libitum* access to food and water. Mice were used at the age of 6-9 weeks. The mice were randomly assigned to the experimental groups and included minimum of 5 mice/group.

6-9 weeks-old wild-type male C57BL/6 were anesthetized by inhalation of 5% isoflurane for 5 min and oropharyngeally administered with 1×10^6^ CFU of T4 strain at mid-log phase, resuspended in 50 μL of 1X endotoxin-free PBS (Invitrogen). Mice were randomly allotted to treatment and control groups (≥5 mice/group). Briefly, 10 mg/kg of SFN in 55% PEG-400 (sigma-Aldrich) and 20% DMSO (HiMedia) was administered by tail vein injection post 1 h of infection and intraperitoneally every 24 h till the clinical endpoint was attained. Mice treated with PEG-400 and DMSO without any SFN served as the placebo control. Animals were clinically scored every 24 h for the symptoms and were sacrificed up on reaching the clinical end points. Euthanasia was done by inhalation of 10% of CO_2_ and cardiac puncture was performed post euthanasia to collect the blood sample. Post sacrifice, cardiac perfusion was performed using endotoxin-free PBS supplemented with 5 mM EDTA (HiMedia) to drain out the blood. Lungs were collected in PBS, homogenized, and strained using 70 μm cell strainers (HiMedia) to a final volume of 5 mL. 100 μL of the lung tissue homogenate was serially diluted and plated of blood agar. Colonies were counted post 16 h of incubation at 37°C and 5% CO_2_.

### Statistical analysis

Data were statistically analyzed using GraphPad Prism v10.4.2. Data represent mean ± SEM. Experiments were performed with three biological replicates. The exact number of biological replicates (N) are mentioned in the respective figure legends. Normality was calculated using the Shapiro-Wilk test and used for the statistical tests. Pairwise comparison of normalized data was analyzed using paired/unpaired t-tests mentioned respectively in the figure legends. Differences were considered significant at ∗, p < 0.05; ∗∗, p < 0.01; ∗∗∗, p < 0.001; ∗∗∗∗,p < 0.0001; ns denotes not significant.

