## Supplemental text for "Sorafenib, a clinically approved kinase inhibitor attenuates *Streptococcus pneumoniae* pathogenesis *in vivo* by targeting serine/threonine kinase StkP"

### Supplementary methods and data

### Supplementary figures

**FIGURE S1: Structure modelling and validation of pneumococcal StkP-kinase domain** (Related to Figure 1) **(A).** Structure of full-length StkP predicted by AlphaFold showing the functional domains. **(B).** Modelled structure of the kinase domain of StkP (StkP-KD) showing catalytically active regions. **(C).** Ramachandran plot of StkP-KD generated using the structure validation tool of SWISS-MODEL, a homology modelling server, **(D).** Sequence and **(E)** structural alignment diagram showing the conservation of the ATP-binding motif, P-loop and catalytically active, C-loop of StkP-KD of *S. pneumoniae* with homologues in *S. aureus (*PknB*)* and *M. tuberculosis (*PknA, PknB*)*, respectively. The PDB ID and RMSD of alignment are highlighted. Zoomed in region shows the structural similarity of catalytic and P-loop regions showing a good structural overlap.

**FIGURE S2: P-loop and C-loop conservation in bacterial and human kinases.** (Related to Figure 1) **(A).** Sequence and (**B).** structural alignment diagrams showing the conservation of P-loop and Catalytic loop (C-loop) of pneumococcal eukaryotic type Ser/Thr kinase domain, StkP-KD and human kinases with available PDB structure, that showed the highest similarity. Microtubule affinity-regulating kinase 3 (MARK3; 7P1L), polo-like kinase 4 (PLK4; 4JXF), AMP-activated protein kinase catalytic subunit alpha-2 (AMPK2; 2H6D), Maternal embryonic leucine zipper kinase (MELK; 4UMT), Salt-Inducible Kinase 3 (SIK3; 8R4O). **(C)**. The C-loop and P-loop regions are magnified to show structural overlap. **(D).** Growth kinetics of *S. pneumoniae* TIGR4 strain in the presence of the top two kinase inhibitor compounds from docking studies, Sorafenib and Dasatinib at 1 μM and 10 μM doses. **** indicates $p \leq 0.0001$ and ns denotes non-significance with respect to equivalent DMSO (solvent) concentrations by Mann-Whitney test. Data is representative of two independent experiments.

**FIGURE S3: Interaction of sorafenib with StkP-KD and StkP-KD R135A/D136A.** (Related to Figure 1) **(A).** Interaction fraction graph showing the time fraction of hydrogen bonds, water bridges and hydrophobic interactions between sorafenib (SFN) and the corresponding amino acids of StkP-KD during a molecular dynamic simulation for 200 ns. Catalytic loop is highlighted, and the active site residues are shown in brown. **(B).** Timeline graph showing the quantification of time-dependent contacts between SFN and StkP-KD during the simulation. Active site residues are represented in brown color. **(C).** Contact diagram showing interactions between SFN and StkP-KD R135A/D136A double mutant. Residues involved in bonding over 30% of the total simulation time are shown. **(D).** Interaction fraction graph showing the time fraction of hydrogen bonds, water bridges and hydrophobic interactions between SFN and StkP-KD R135A/D136A. The C-loop region is highlighted, and the mutated residues are shown in brown. **(E).** Timeline graph showing the quantification of time-dependent contacts between SFN and StkP-KD R135A/D136A during the simulation.

**FIGURE S4: Competitive binding of Sorafenib and DivIVA peptide with StkP-KD** (Related to Figure 1) **(A).** Contact diagram of interactions between sorafenib (SFN) and StkP-KD, during a 200 ns molecular dynamic simulation in the presence of DivIVA peptide. Residues involved in bonding over 30% of the total simulation time are shown. (**B**). Interaction fraction graph showing the time fraction of hydrogen bonds, water bridges and hydrophobic interactions of SFN with StkP-KD in the presence of DivIVA peptide. Catalytic loop is highlighted, and the active site residues are shown in brown. **(C).** Contact diagram showing the interactions between DivIVA peptide and StkP-KD during the simulation in the presence of SFN. Only residues involved in bonding over 30% of the total simulation time are shown. **(D).** Interaction fraction graph showing the time fraction of interactions between DivIVA peptide and StkP-KD during the simulation in the presence of SFN. Catalytic loop is highlighted, and the active site residues are shown in brown.

**FIGURE S5: Validation of D39-gfp-stkP($P_{Zn}$) and D39ΔstkP::gfp-stkP($P_{Zn}$) strains** (Related to Figure 2) **(A).** Flow cytometry overlay showing the positive shift in GFP signal in D39-gfp-stkP($P_{Zn}$) strain upon induction with 0.5 mM $ZnCl_2$ and 0.05 mM $MnCl_2$ compared to uninduced D39-gfp-stkP($P_{Zn}$) and D39-WT strain. Percentage positivity is shown in parenthesis. **(B).** Western blot with anti-GFP showing the dose-dependent increase in expression of GFP-StkP in D39-gfp-stkP($P_{Zn}$) with increasing concentrations of $Zn^{2+}$ and $Mn^{2+}$ (10:1). **(C).** PCR amplification of StkP gene fragment in D39-WT, D39ΔstkP::gfp-stkP($P_{Zn}$) (complemented strain) and D39-gfp-stkP($P_{Zn}$) (overexpression strain). Equal amount of genomic DNA was used as the template. **(D).** Flow cytometry overlay and (**E**) confocal microscopy images showing the GFP-StkP expression and septal localization in D39ΔstkP::gfp-stkP($P_{Zn}$) upon induction with 0.5 mM $Zn^{2+}$ and 0.05 mM $Mn^{2+}$ compared to uninduced D39ΔstkP::gfp-stkP($P_{Zn}$) and D39-WT strain. Inset shows the magnification of selected region. Scale bars, 5 µm. **(F, G).** Growth kinetics of **(F).** D39-gfp-stkP($P_{Zn}$) and **(G).** D39ΔstkP::gfp-stkP($P_{Zn}$) in the presence of 0.5 and 1 µM DMSO under uninduced and induced conditions. ns denotes no significance by Mann-Whitney test comparing uninduced/induced

DMSO treatments to respective untreated bacteria. Data in panels F, G are representative of mean ± SEM from three independent experiments.

**FIGURE S6: Effects of sorafenib on StkP mediated phosphorylation** (Related to Figure 3) **(A-D).** Uncropped western blots probed with pThr specific antibody showing the downregulation of phosphorylated proteins in (**A**) D39 and (**C**) TIGR4 (T4) strains upon treatment with 10 μM sorafenib (SFN). Untreated (UTD) and equivalent amount of DMSO treated bacteria were used as the controls. Labelled bands showed consistent downregulations in both strains. (**B, D**). Enolase was used as the loading control. **(E).** Coomassie stained SDS-PAGE and **(F).** western blot with anti-His antibody of the purified recombinant 6x-His tagged StkP-KD protein. **(G-J).** Western blots showing the *in vitro* autophosphorylation of purified StkP-KD **(G).** under a gradient of ATP concentrations (0-500 μM) and **(I).** upon treatment with equimolar and twice concentration of SFN relative to ATP (250 μM). **(H, J).** Ponceau staining of the blots in panels G and I respectively, were used as loading control. **(K).** Densitometry analysis of the blot in panel I normalized to total protein. Data in G-K are representative of two independent experiments. **(L).** Coomassie stained SDS-PAGE of total cell lysate (20 μg) of *E. coli* BL21(DE3) transformed with KDNC2 plasmid (StkP-KD) and treated with DMSO and SFN respectively. pET-22b(+) empty vector transformed (20 μg) and KDNC2 induced bacteria (2 μg) served as controls. **(M).** Growth kinetic assay showing the growth of *E. coli* BL21(DE3) cells transformed with pET-22b(+) vector and KDNC2 plasmid upon treatment with 10 μM and 50 μM SFN for 5 h. ns denotes non-significance of SFN relative to DMSO concentration by Mann-Whitney test.

**FIGURE S7: Antimicrobial effect of sorafenib with limited host cytotoxicity** (Related to Figure 4) **(A).** PrestoBlue cell viability assay showing the cytotoxicity of A549 cells under 2.5 to 10 μM of sorafenib (SFN) and DMSO for 3 h. 5 μM Staurosporine treated cells were used as the positive control for cell death. **(B).** Flow cytometry analysis showing the quantification of propidium iodide positive *S. pneumoniae* T4 upon treatment with 10 μM SFN. 70% isopropanol treated bacteria were used as the positive control of death. **(C).** CFU plating assay showing the killing of 10 μM SFN treated T4 strain upon incubation with 10% normal human serum (NHS). Bacteria treated with equimolar DMSO concentration served as a control. Bacteria incubated with heat-inactivated serum was used as the negative control. * indicates $p \leq 0.05$, ** indicates $p < 0.01$ and ns denotes non-significance by paired t-test. Data represents mean ± SEM from three independent experiments.

### Supplementary tables

**Table S1:** (Related to Figure 1 and S2). Kinase inhibitors used for virtual screening against pneumococcal StkP-KD, their CAS and PubChem IDs. Retrieved from (Llovet *et al*, 2008).

**Table S2:** (Related to Figure 1 and S2). Docking scores of compounds (kJ/mol) with StkP-KD from virtual screening and their PubChem IDs. Multiple entries of the same compound indicate alternative docked confirmations. Second sheet contains the raw data file from Schrodinger.

### Supplementary video legends

**Video S1:** Movie showing the interaction of sorafenib with the active site residues of pneumococcal StkP-KD. Sorafenib is shown as a ball and stick model with elemental colorings, Carbon in white, Oxygen in red, Nitrogen in blue, Fluorine in light green and Chlorine in dark green. StkP-KD backbone is shown as a cartoon in green color with the functionally active domains P-loop (19-25) in blue, C-loop (133-141) in red, activation loop (162-173) in purple and P+1-loop (174-179) in yellow colors respectively. The active site residues Arg-135 and Asp-136 are also represented as ball and stick model in cyan and pink colors respectively. Yellow dotted lines indicate the hydrogen bonds formed between sorafenib and catalytic residues. Movie was made using the Maestro module of Schrodinger Software Suite.

**Video S2:** Zoomed view of the catalytic cleft in video S1.

**Video S3:** Movie showing the drifting of sorafenib away from the catalytic cleft upon mutating the catalytic residues, Arg-135 and Asp-136 to alanine. Color and model representations are the same as followed in movie S1. Mutated residues Ala-135 and Ala-136 are shown in cyan and pink colors respectively.

**Video S4:** Interaction of sorafenib with the catalytic residue Arg-136 (cyan color) in the presence of DivIVA peptide. The peptide is shown as a stick model in orange color and Ser-165 is shown in lavender color. Rest all the representations are same as video S1.

**Video S5:** Movie in S4 zoomed to the catalytic cleft of StkP-KD.

### Supplementary Methods

**Protein preparation for docking**

The predicted model of the StkP Kinase Domain was imported into the Maestro software [Schrödinger Release 2023-2: Maestro, Schrödinger, New York,]. The model was prepared using the Protein Preparation Wizard (Sastry *et al*, 2013) [Schrödinger Release 2024-4], which involved preprocessing to add hydrogen atoms, assigning bond orders, and creating zero-order bonds to metals and disulfide bonds. Water molecules were deleted unless they were involved in critical interactions. Subsequently, hydrogen bond assignments were optimized to improve the overall structural stability. Finally, the structure underwent energy minimization to resolve any steric clashes and to refine the geometry, ensuring the model was suitable for subsequent molecular modelling studies.

**Ligand preparation**

The individual structures of mammalian kinase inhibitors (**Table S1**) were retrieved from the PubChem database (https://pubchem.ncbi.nlm.nih.gov/) and prepared using the LigPrep module [Schrödinger Release 2024-4: LigPrep]. The preparation followed the default settings of the tool. Briefly, the structures were processed using the OPLS4 (Optimized Potentials for Liquid Simulations) force field (Lu *et al*, 2021). Possible ionization states at a target pH of 7.0 ± 2.0 were generated using the Epik (Classic) program (Johnston *et al*, 2023) [Schrödinger Release 2024-4]. The process excluded consideration of metal binding states and the original ionization state. Additionally, the structures were desalted, and tautomers were generated, with the ligand size limited to a maximum of 500 atoms. Stereoisomers were generated while retaining specified chiralities, with a limit of 32 structures per ligand. The prepared ligand structures were saved in Maestro format.

**Receptor grid generation**

A binding pocket identified by the SiteMap tool (Halgren, 2009) [Schrödinger Release 2024-4], containing the active site residue Asp 136 along with other catalytic loops, was selected for grid generation for docking studies. The grid was generated using the Receptor Grid Generation tool of Schrödinger, following default settings. The Van der Waals radius scaling used a scaling factor of 1.0 and a partial charge cutoff of 0.25, without applying any per-atom scaling factors or considering aromatic hydrogen and halogen hydrogen bonds. The grid was generated at the site defined by the SiteMap tool, without incorporating any constraints, rotatable groups, or excluding volumes.

**Cloning and purification of StkP kinase domain**

pET22b(+) vector was double digested using Hind-III and Nde-I (GeneI) restriction enzymes following GeneI protocol: incubation at $37^{o}C$ for 4 hrs, followed by heat inactivation at $65^{o}C$

for 20 min. The digested product was run in 1% agarose gel and required band was isolated by gel extraction using MN kit.1-277 amino acid region corresponding to the nucleotide of the CDS of StkP - Kinase Domain (NCBI: NC_003028.3) was amplified using the following primers at an annealing temperature of 55ºC.

Forward: GATATACATATGatccaaatcggcaagatt

Reverse: GGCCGCAAGCTTacgattgtaggacaagctacta

The PCR products were purified by PCR cleanup and then were digested using Hind-III and Nde-I, followed by clean up. Digested product and the vector were ligated, in the insert to vector molar ratio 15:1, using T4 DNA ligase by incubating at 16ºC for 3 hrs. The ligated product was transformed into *E. coli* Mach1 cells. Colonies were screened by colony PCR using the same set of primers. Plasmids were isolated from positive colonies, and the clone was confirmed by double digestion as well as Sanger sequencing. The product consists of the C-terminal 6x-His tag region followed by 1-277 residues of pneumococcal StkP- Kinase Domain (KDNC2 plasmid).

For protein expression, the plasmid construct was transformed into *E. coli* BL21 (DE3) cells and plated onto an LB agar plate with 100 µg/mL of ampicillin. A few colonies from the transformation plate were seeded into primary culture in 15 mL 2xYT medium supplemented with 100 µg/mL of ampicillin for 12 h at 37°C and 200 rpm. The primary culture was inoculated into 1 L of 2xYT medium as a secondary culture and allowed to grow at 37°C in a shaker incubator at 200 rpm until the $OD_{600}$ value reached 0.5. The culture was then induced with 0.5 mM IPTG and allowed to express protein at 37°C for 4 hrs. The bacterial cells were harvested by centrifugation, and the cell pellet was resuspended in a lysis buffer containing 50 mM Tris-HCl (pH 7.5), 500 mM NaCl, 10 mM imidazole, 1 mM Phenylmethylsulfonyl fluoride (PMSF), 1 mM benzamidine hydrochloride, and 3 mM β-mercaptoethanol, followed by sonication for 8 min at 40% amplitude with 2 s on and 5 s off cycles. The soluble fraction was collected by a high-speed spin at 13,000g for 30 min at 4°C. The supernatant was filtered using a 0.45 µm syringe filter. The filtered supernatant was used for the affinity chromatography.

For affinity purification of the protein, an EconoFit Nuvia IMAC Ni-NTA column of 5 mL volume (Bio-Rad) connected to an NGC Quest 10 protein purification system (Bio-Rad) was used. The column was pre-equilibrated with 8x column volume of the lysis buffer, followed by sample injection. The column was then washed with 8x column volume of a wash buffer [20 mM Tris-HCl (pH 7.5), 300 mM NaCl, 25 mM imidazole, 1 mM PMSF, 1 mM benzamidine hydrochloride, and 3 mM β-mercaptoethanol], followed by elution with 10x column volume of an elution buffer containing 400 mM imidazole. The peak fractions were loaded on a 12% SDS-PAGE, and the fractions containing the protein were combined and concentrated to 5 mL using a Vivaspin centrifugal concentrator with 10,000 MWCO (Sartorius). The concentrated sample was given a high-speed spin at 13,000g for 30 min at 4°C to remove the precipitates. The collected supernatant was used for the size-exclusion chromatography step using a HiPrep16/60 Sephacryl HR S-200 column (Cytiva), pre-equilibrated with 1x column volume of a size-exclusion buffer containing 20 mM Tris- HCl (pH 7.5), 300 mM NaCl, 1 mM EDTA, and 1 mM DTT, and connected to an NGC Quest 10 protein purification system. The peak fractions were analyzed on a 12% SDS-PAGE, and the pure fractions were pooled together and concentrated to 1 mL. The concentrated protein sample was dialyzed into 1x PBS and was aliquoted, flash-frozen in liquid nitrogen, and stored in -80°C. The purity of the final protein was checked by running on a 12% SDS-PAGE and Western blotting with anti-His antibody (Sigma; 1: 2000).

***In vitro* autophosphorylation assay**

Phosphorylation of the purified StkP-KD was performed *in vitro* following the protocols by (Stauberova *et al*, 2024) and (Li *et al*, 2022) with minor modifications. Briefly 1 µM of purified StkP-KD was added to 30 µL reaction buffer containing 25 mM Tris-HCl (pH 7.4), 100 mM NaCl, 1 mM DTT, 0.1 mM EDTA and 10 mM $MnCl_2$. The reaction was started by adding ATP and was incubated at 37ºC for 30 min. The reaction was stopped by 10 µL of 4X Lamelli sample buffer (Bio-Rad) supplemented with 50 mM DTT (HiMedia) followed by boiling at 70ºC for 10 min. The samples were resolved on a 12% SDS-PAGE and western blotting with anti-Phospho-Threonine monoclonal antibody (CST) was performed as described earlier. The membrane post transfer was stained with Ponceau (Invitrogen) and was used as the loading control. Initial set of experiments were performed with a range of ATP concentrations to identify the concentration at which an increase in the basal phosphorylation was occurring. For assays involving sorafenib, the drug was added to the mixture at 1:1 and 1:2 molar ratio relative to ATP concentration and the kinase reaction was initiated was initiated by ATP addition.

**PrestoBlue cytotoxicity assay**

A549 cells were seeded in a 96-well plate (seeding density 10,000 cells/well) and incubated overnight at 37°C and 5% $CO_2$. The media was removed and 100 µL DMEM supplemented with 10% FBS and 1% penicillin-streptomycin was added. The cells were treated with sorafenib and DMSO at range of concentrations between 2.5-10 μM and incubated for 3 h. 5 μM staurosporine was used as a positive control to induce cell death by apoptosis. After the incubation time, the media was removed, and two PBS washes were given. Fresh 90 μL DMEM complete media was added along with 10 μL of PrestoBlue HS reagent (Invitrogen) and incubated for 10 min. The fluorescence was measured in a Varioskan LUX multimode microplate reader (ThermoFisher) at 560 nm excitation / 590 nm emission. The percentage viability was calculated by normalizing to the untreated control.

**Flow cytometry of bacterial viability by Propidium Iodide staining**

Propidium Iodide (PI) staining for quantifying dead bacterial cells was performed following the live/dead staining assay protocol with minor modifications. Briefly, ~$5 \times 10^7$ bacterial cells were harvested post-treatment with either sorafenib or DMSO, washed once with PBS, and resuspended in 500 μL PBS. To this, 1.5 μL of PI dye was added, and the suspension was incubated in the dark for 20 min at room temperature. Following incubation, cells were washed twice with PBS and fixed using 4% paraformaldehyde (PFA, HiMedia) for 10 min at room temperature. Fixed cells were centrifuged and resuspended in PBS for flow cytometric analysis. As a positive control for cell death, bacteria were treated with 70% isopropanol for 20 min at room temperature with shaking at 150 rpm prior to staining. Flow cytometry was performed using a CytoFLEX S Flow Cytometer (Beckman-Coulter), detecting PI-positive cells in the ECD channel (Texas Red).
