## Supplementary figures and images for "Sorafenib, a clinically approved kinase inhibitor attenuates *Streptococcus pneumoniae* pathogenesis *in vivo* by targeting serine/threonine kinase StkP"

### Figure S1

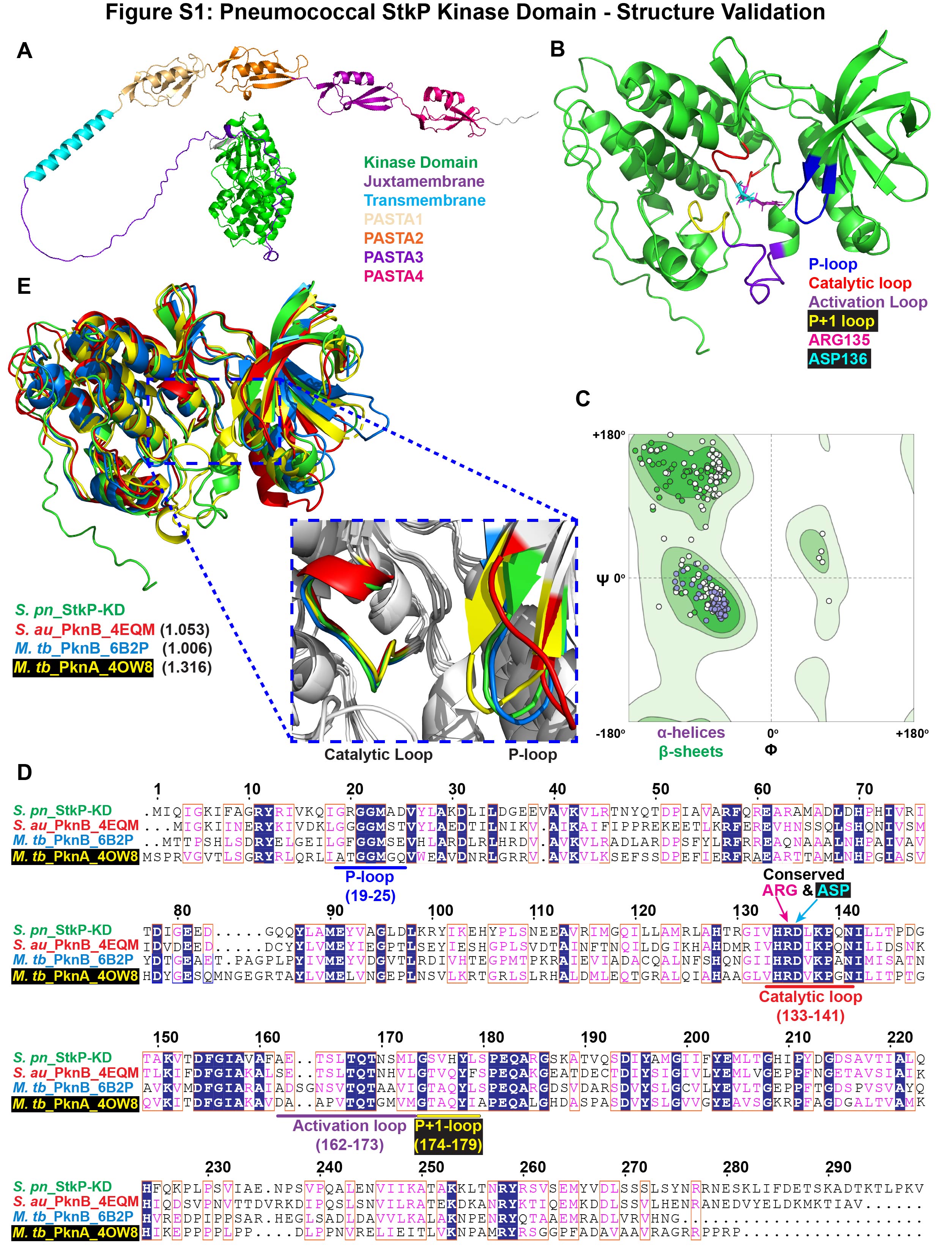

### Figure S2

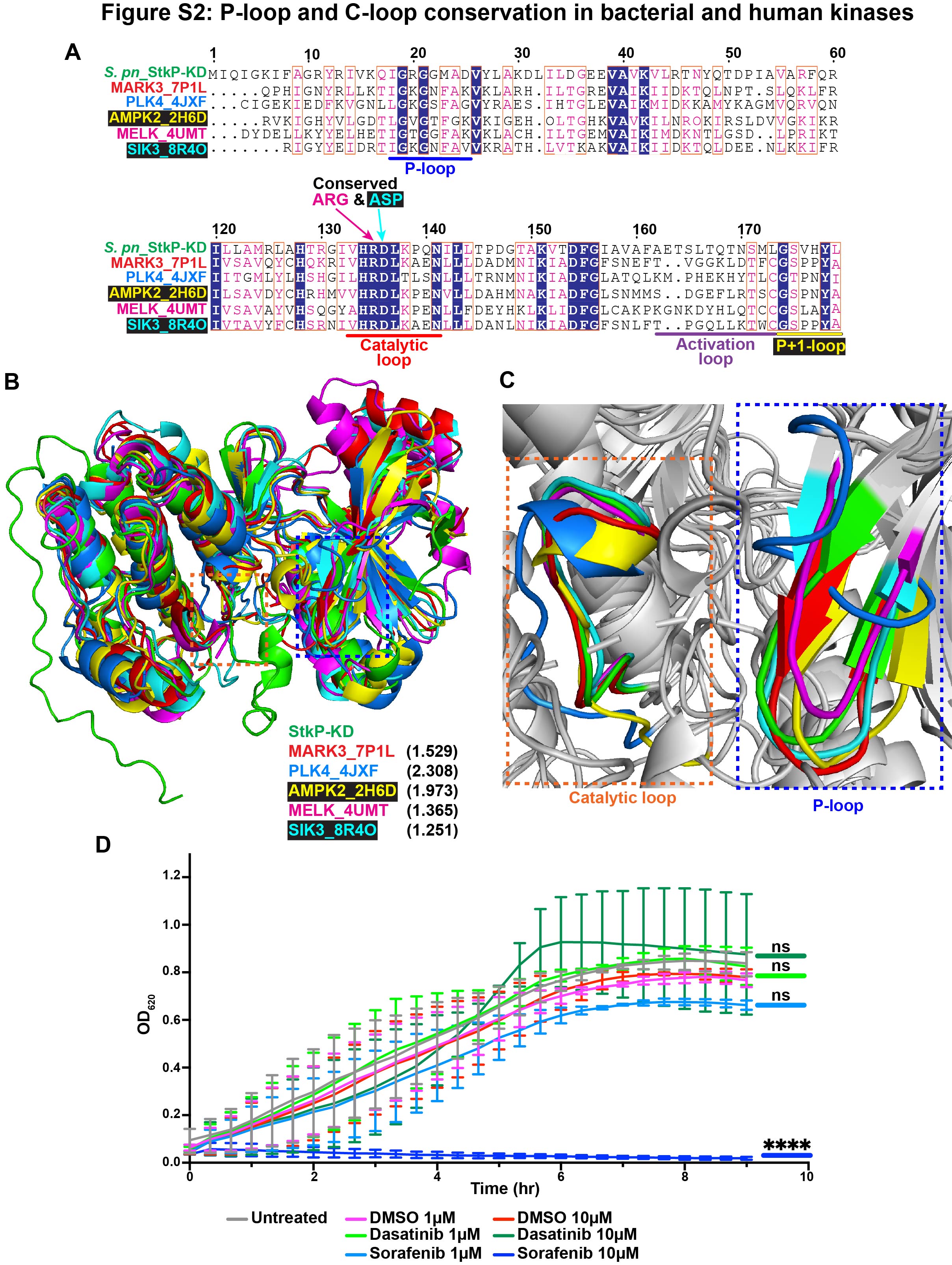

### Figure S3

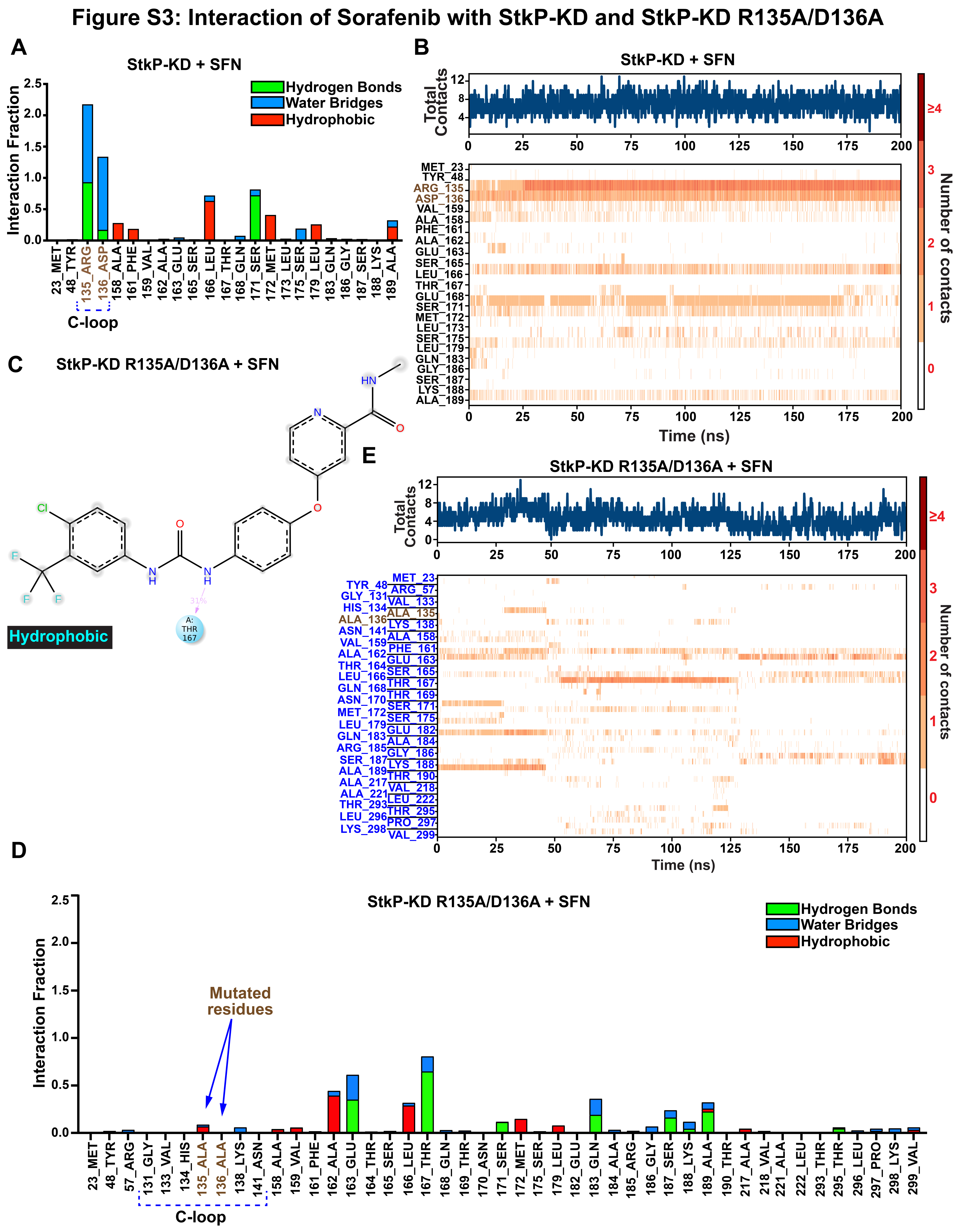

### Figure S4

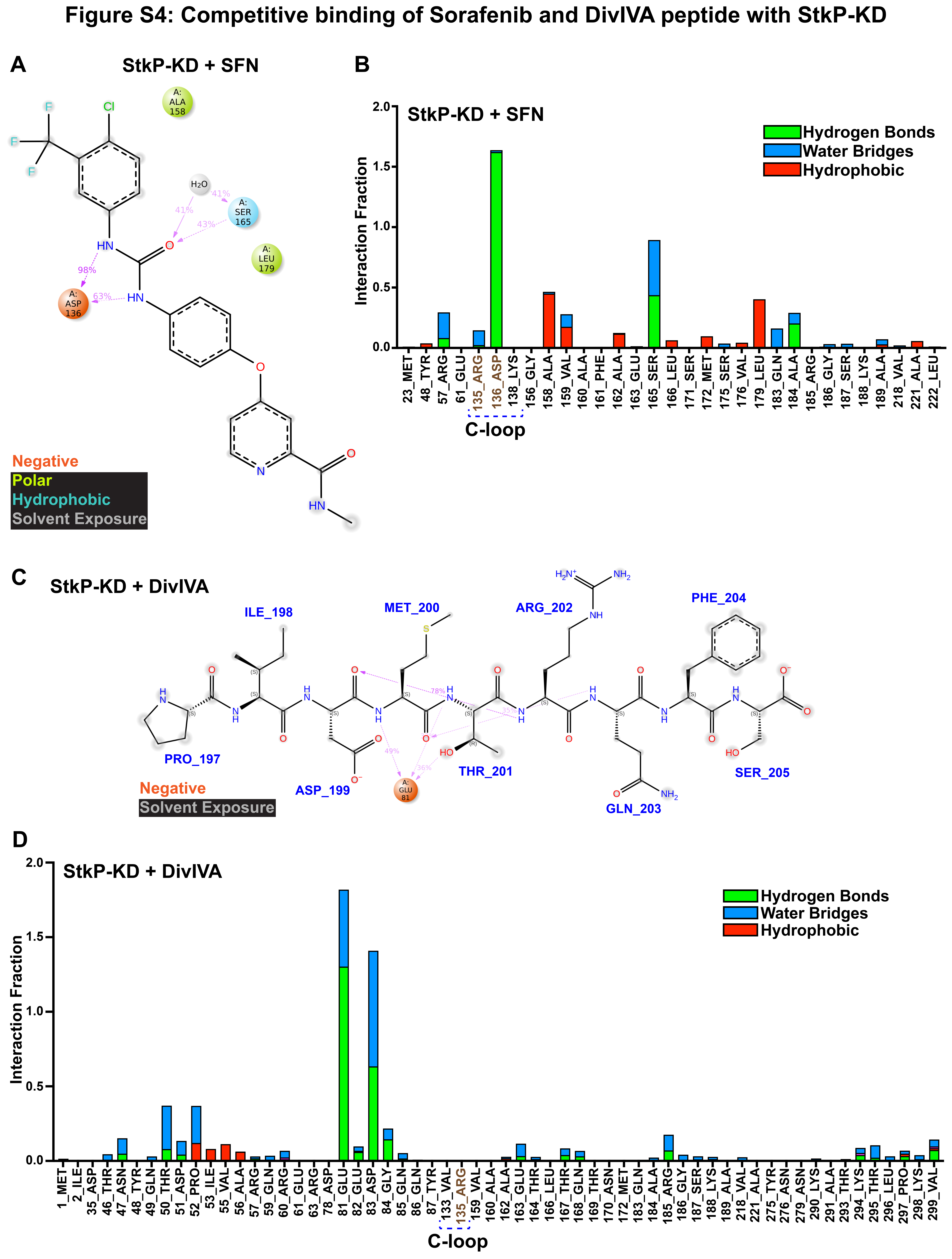

### Figure S5

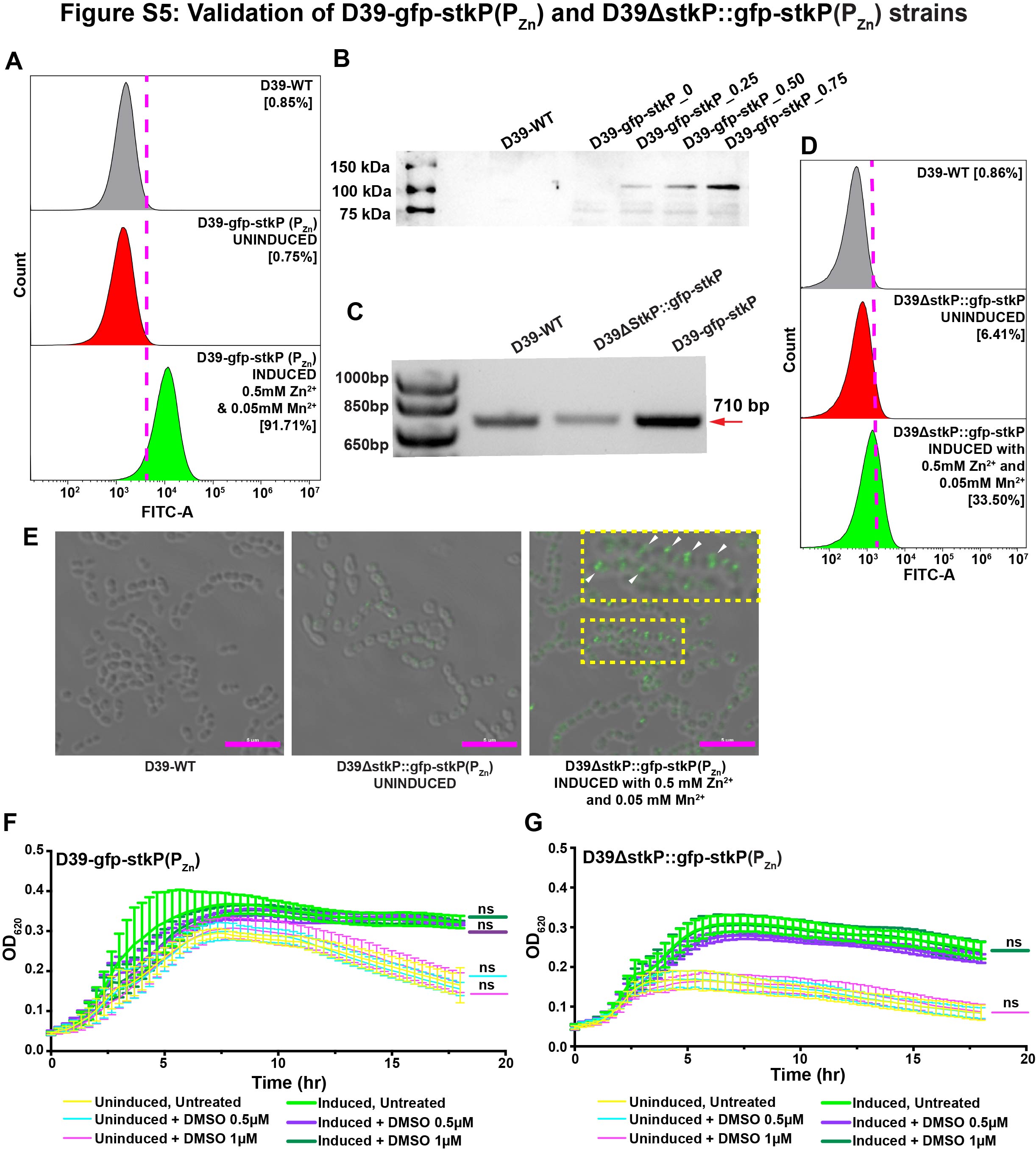

### Figure S6

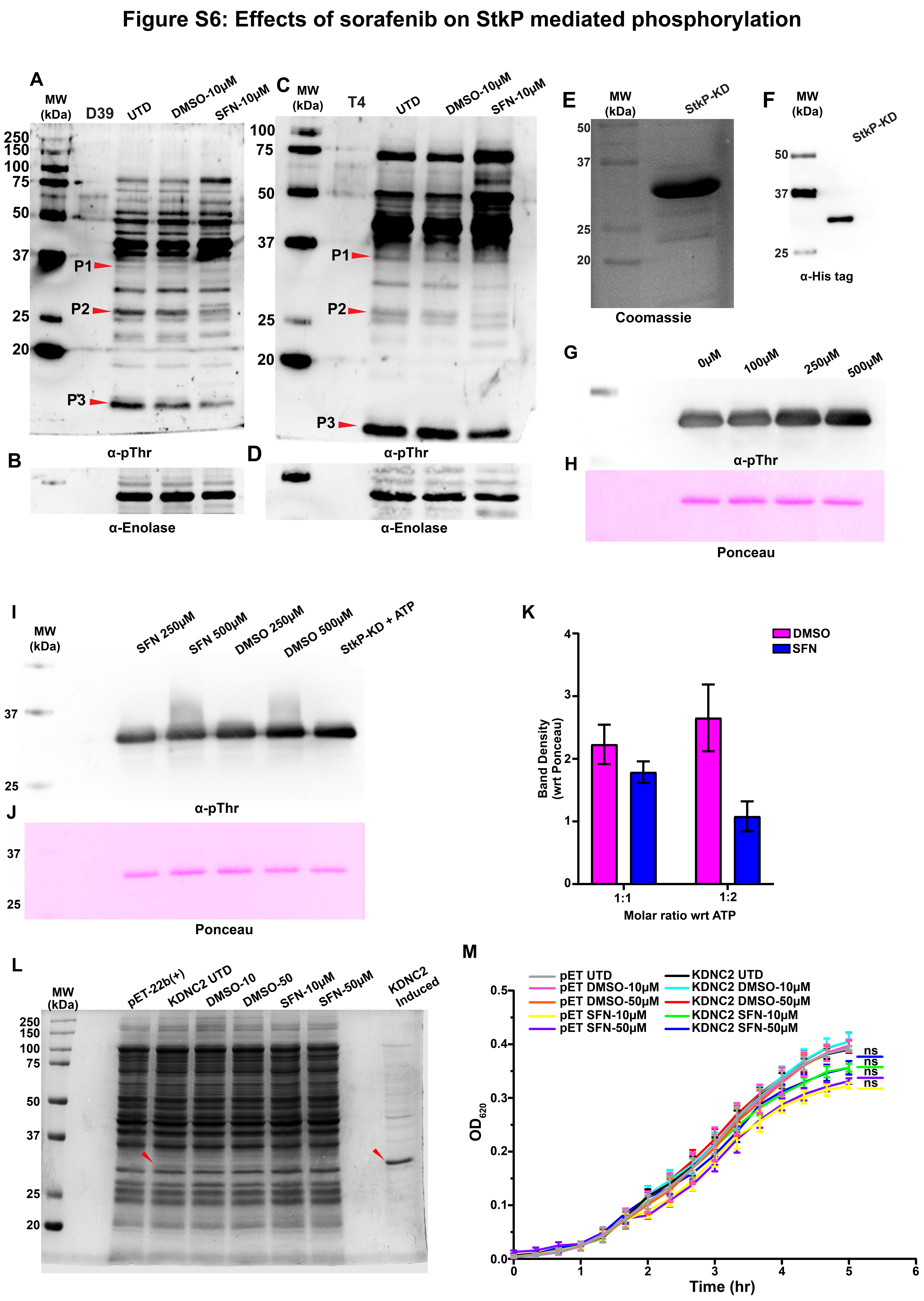

### Figure S7

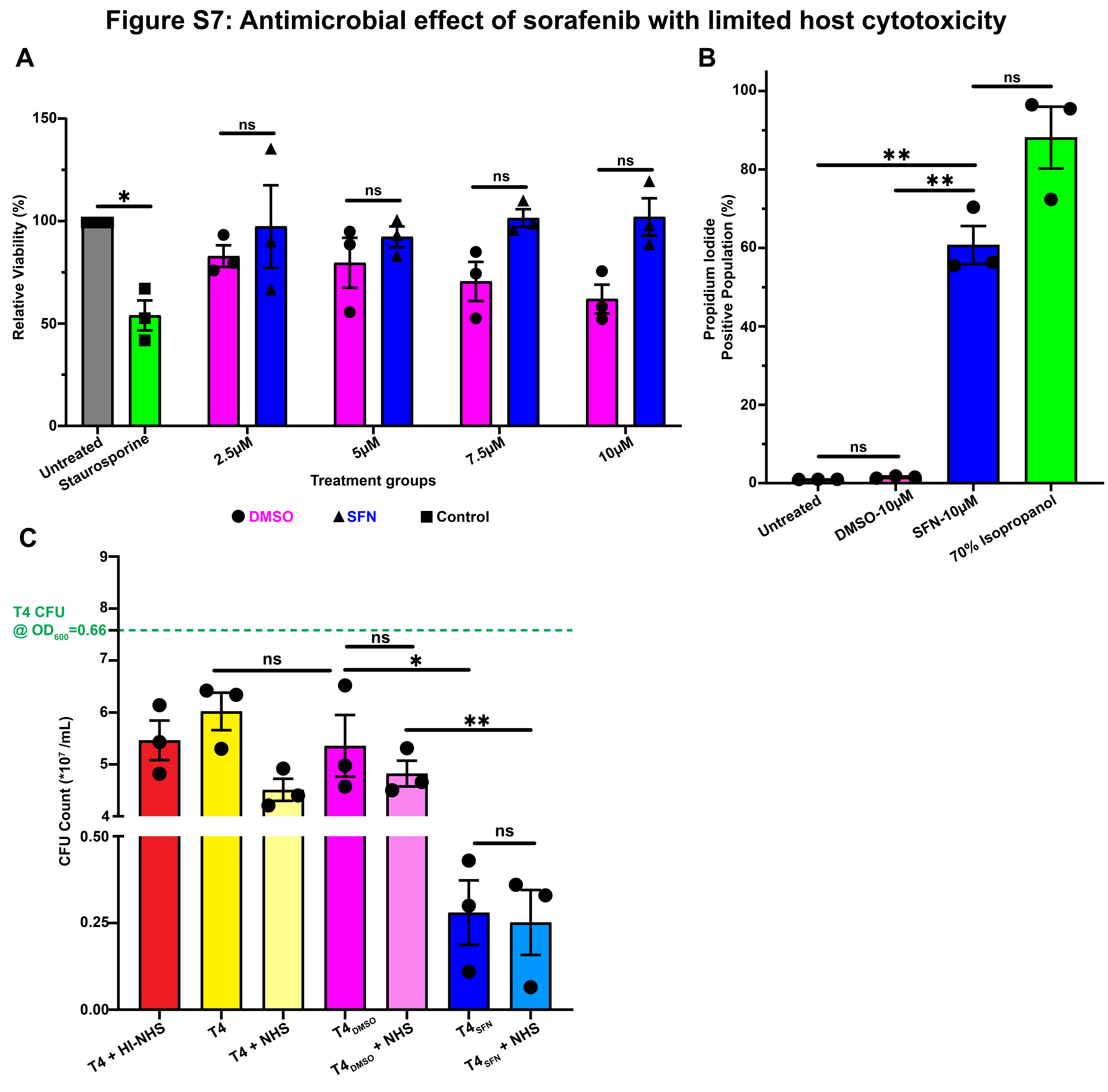
